# Uncovering synaptic and cellular nanoarchitecture of brain tissue via seamless *in situ* trimming and milling for cryo-electron tomography

**DOI:** 10.1101/2025.03.09.642162

**Authors:** Jiying Ning, Jill R. Glausier, Rangana Warshamanage, Leslie Gunther-Cummins, Tom Burnley, Colin M. Palmer, Guillermo Gonzalez-Burgos, Takeaki Miyamae, Jing Wang, Diane Carlisle, Chyongere Hsieh, Thomas Schmelzer, Silas A. Buck, Jonathan Franks, Cheri M. Hampton, William R. Stauffer, David A. Lewis, Robert M. Friedlander, Frank P. Macaluso, Martyn Winn, Michael Marko, Zachary Freyberg

## Abstract

Cell-cell communication underlies all emergent properties of the brain, including cognition, learning and memory. The physical basis for these communications is the synapse, a multi-component structure requiring coordinated interactions between diverse cell types. However, many aspects of three-dimensional (3D) synaptic organization remain poorly understood. Here, we developed an approach, seamless *in situ* trimming and milling (SISTM), to reliably fabricate sufficiently thin lamellae for mapping of the 3D nanoarchitecture of synapses in mouse, monkey and human brain tissue under near-native conditions via cryo-electron tomography (cryo-ET). We validated SISTM in a mouse model of Huntington’s disease, demonstrating distinct 3D alterations to synaptic vesicles and mitochondria. By successfully applying SISTM to macaque brain, we described the 3D architecture of a tripartite synapse within the cortex. Subtomogram averaging (STA) enabled spatial mapping of astrocyte-neuron contacts within the tripartite synapse, revealing neurexin-neuroligin complexes as potential constituents that tether the two cell types. Finally, we showed that the defining features of synaptic nanoarchitecture were conserved across species and evident in human brain tissue obtained postmortem. Combining SISTM with cryo-ET and STA is a starting point for a new understanding of brain organization, disease-induced structural alterations and the development of rational, structure-guided therapeutics.

## Main

The cryo-electron microscopy (cryo-EM) revolution has enabled the visualization of macromolecules at atomic resolution. Indeed, cryo-EM single-particle analysis (SPA) is the ascendant approach for high-resolution structural reconstructions of vitrified macromolecules^1,2^. However, most SPA samples are purified and therefore imaged *in vitro* outside of their native cellular environments. The development of *in situ* cryo-imaging approaches that visualize structures in their native states within cells, and in the absence of chemical fixation or other extrinsic manipulations that potentially distort cytoarchitecture, promises many important biological insights^3^. The advent of *in situ* cryo-electron tomography (cryo-ET) has begun delivering on this promise by three-dimensional (3D) imaging of macromolecules and organelles directly within cells in healthy and disease states^4–7^.

The application of *in situ* cryo-ET to the imaging of intact mammalian tissue represents the next frontier of structural cell biology. A core principle of *in vivo* physiology is the coordinated activity of different cell populations within the highly interactive tissue milieu. The brain best exemplifies this principle given its highly complex, heterogenous cellular makeup^8–12^. Indeed, neurons exhibit distinct, highly polarized cytoarchitectures that include morphologically complex pre- and postsynaptic processes^13–15^. Individual neurons may receive upwards of 30,000 synaptic contacts from other neurons, which represent the basic units of neuronal communication^16–18^. These contacts are highly plastic, and make adaptive changes based on their local environments^13,15,19^. Within brain tissue, neurons also require non-neuronal glial cells, such as astrocytes and oligodendrocytes, to maintain synaptic communication^20,21^. Furthermore, neuropsychiatric diseases are closely associated with disruptions of these complex connections^22–28^. Thus, the ability of *in situ* cryo-ET to directly image neurons, including their sub-cellular compartments and connections, directly in brain tissue can provide novel insights into brain function and dysfunction.

Major technical challenges persist in applying cryo-ET to tissues, and brain in particular. Foremost, the thickness of intact tissues makes it difficult to visualize subcellular details due to inadequate penetration of the electron beam. To address this limitation, cryo-focused ion beam (cryo-FIB) milling has been increasingly employed to generate lamellae sufficiently thin for cryo-ET, particularly for imaging of isolated cells^29–32^. Given the greater thickness of vitrified tissue specimens compared to individual cells, the application of cryo-ET to tissues has been much more limited^33–38^. This paucity of tissue-based studies is based on several factors including challenging tissue preparation conditions since fresh, unfixed tissue is soft and difficult to handle. Another key challenge is production of sufficiently thin tissue samples for cryo-ET. The “lift-out” technique has attempted to address this challenge by enabling extraction of thin lamellae for cryo-ET^39^. Yet, despite recent improvements, the lift-out approach remains technically challenging with extremely low success rates for most users, making it difficult to adapt to high-throughput applications^39,40^.

Due to the above technical challenges, most brain-related cryo-ET studies have bypassed brain tissue, relying primarily on *in vitro* neuronal preparations^41^ or synaptosomes^42^, which consist of synaptic contacts isolated from their native tissue contexts. Furthermore, the few published *in situ* cryo-ET studies of intact brain tissue^36,37,43^ were either disease-focused, concentrating on pathological structures in pre-clinical and clinical Alzheimer’s disease (AD), or described tool development. To fully capitalize on the advantages unique to *in situ* cryo-ET in intact brain tissue, we focused on questions concerning synapses and neuron-glia connections because these structures serve as exemplars of complex cell-cell connections essential for brain organization and function. Moreover, intact tissue avoids potential confounds associated with some experimental models such as isolated cultured cells, which fail to capture the complexity of the synaptic neuropil that exists in tissue. These model systems also often lack the capacity to investigate connections between different cell types, such as neurons and astrocytes. Consequently, many critical questions remain, including: 1) What is the structure and organization of synaptic nanoarchitecture *in situ* and how is it impacted by disease states? 2) How do non-neuronal glial processes interact with synapses from ultrastructure to the molecular level? and 3) Are the defining features of *in situ* synaptic architecture conserved across species and evident in human brain tissue obtained postmortem? To begin answering these questions and make the approach more accessible, here we established a workflow termed seamless *in situ* trimming and milling (SISTM) for cryo-ET imaging of brain tissue.

### Design of the SISTM workflow

Our SISTM workflow for *in situ* cryo-ET imaging of mammalian brain tissue under near-native conditions incorporates the following steps: **(i)** high-pressure freezing (HPF) in a modified carrier that reliably vitrifies brain tissue. **(ii)** Exposing and trimming frozen brain tissue via cryo-ultramicrotomy, followed by **(iii)** cryo-FIB-milling of the trimmed tissue to efficiently generate ultrathin lamellae for **(iv)** *in situ* cryo-ET imaging (**Figure 1a**). This approach uses on-carrier milling, avoiding lift-out and offering a more complete tilt range (±60°) (**Figure 1b-g** and **Extended Data Figure 1**). Further advantages of this workflow include greater sample stability due to on-carrier milling, generation of multiple usable lamellae from a single tissue sample, and seamless integration into existing downstream workflows for cryo-transmission electron microscopes (cryo-TEM). Additionally, the SISTM method closely matches the geometry of lamella preparation (*e.g*., milling angle) with on-grid cell milling, which makes it possible to modify the existing autoTEM software template for a semi-automated milling process. This results in a more user-friendly approach compared to lift-out methods which rely on more time-consuming manual operation.

**Figure 1.**
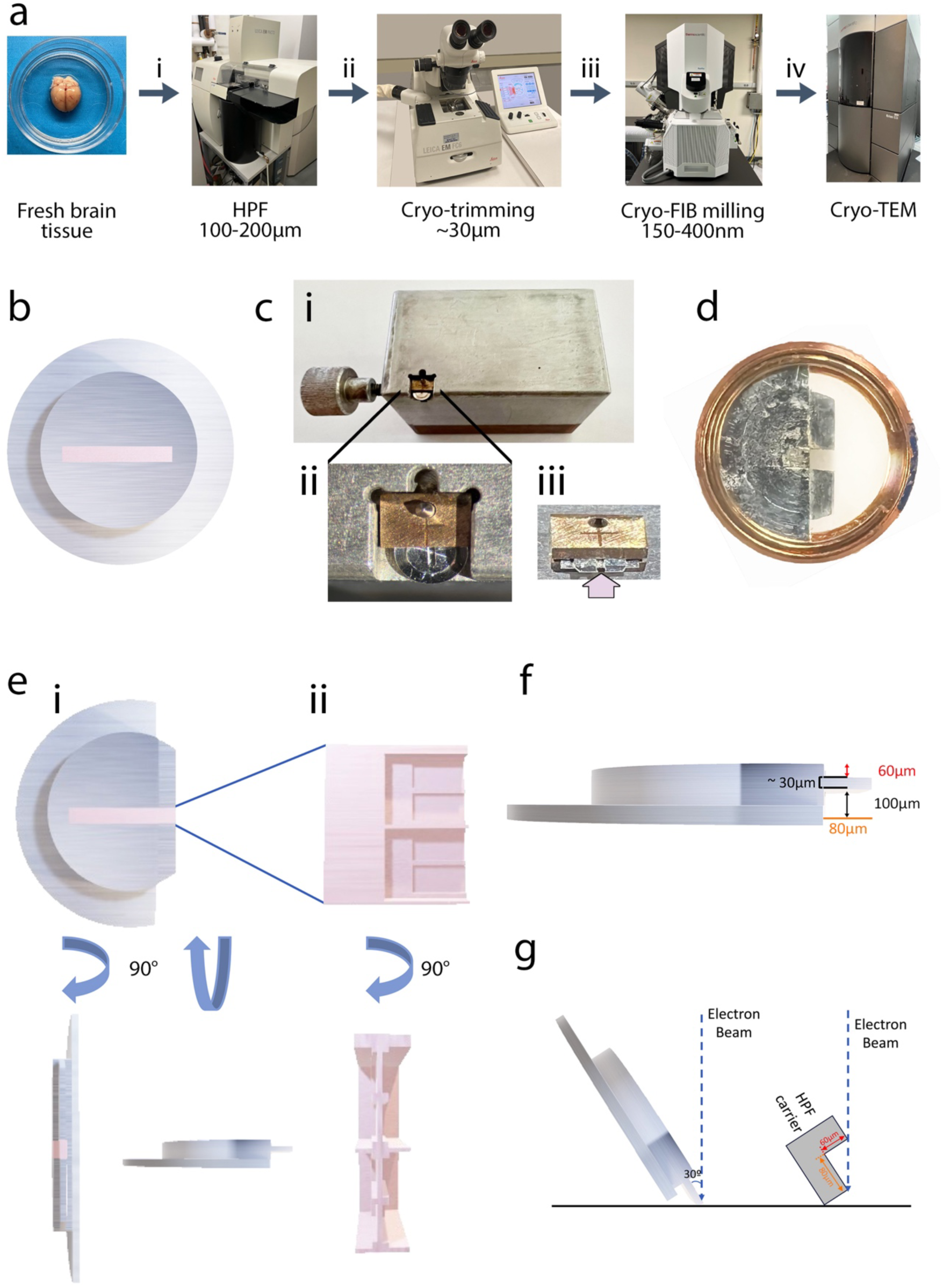
SISTM workflow for preparation of fresh brain tissue for *in situ* cryo-electron tomography. **(a)** Overview of the workflow. (**i**) A 100-200 µm-thick sample of fresh brain tissue is painted into an aluminum carrier which is loaded into a high-pressure freezer (HPF) for immediate cryo-preservation; a representative image of fresh mouse brain is shown. (**ii**) The high-pressure frozen sample is subsequently trimmed in a cryo-ultramicrotome to ∼30 µm thickness. (**iii**) The sample is thinned via cryo-focused ion beam (cryo-FIB) milling to a final 150-400 nm thickness followed by (**iv**) imaging of the lamellae in a cryo-transmission electron microscope (cryo-TEM). **(b)** An overhead schematic of the HPF carrier with brain tissue (pink) filling the carrier slot. **(c)** For cryo-trimming, the aluminum HPF carrier is loaded into an Intermediate Sample Holder (ISH) by means of a loading block (**Panel i**), which releases the spring tension. The HPF carrier is supported during trimming by the spring tension of the ISH (**Panel ii**). A representative image of the cryo-trimmed carrier in the ISH; arrow indicates the sample location (**Panel iii**). **(d)** A representative image of the trimmed HPF carrier clipped into an autogrid ring to facilitate subsequent shuttling of the sample for cryo-FIB milling and imaging. **(e)** Schematic illustrating the progressive thinning of cryo-preserved brain tissue. The HPF carrier is first trimmed away via cryo-ultramicrotomy to expose brain tissue (**Panel i** with 90° views). This is followed by progressive cryo-FIB milling to fabricate lamellae that are sufficiently thin to enable cryo-electron tomography (cryo-ET) imaging (**Panel ii** with 90° view)**. (f)** Overview of the cryo-trimming strategy of the HPF carrier for cryo-ET. Side view of the trimmed carrier illustrates the optimal cryo-trimming of ∼60 μm from the top of the carrier (red double arrow) and ∼100 μm from the bottom of the carrier (black double arrow). This leaves a 20-30 μm-thick shelf of tissue exposed on the top and bottom that is stabilized by the remaining carrier on the sides. The tissue shelf extends ∼80 μm from the body of the carrier (orange). **(g)** When the carrier is oriented at a 60° tilt angle, a minimal trimming depth of ∼80 µm (orange), represented by the tissue shelf, permits unobstructed imaging across the full tilt angle range (±60°).

Brain tissue was placed into custom-designed aluminum carriers (**Figure 1b**) using a fine-tipped brush, followed by immediate vitrification via HPF. A major advantage of our carrier design is the capability to clip the carriers into autogrid rings. This avoids switching between different adaptors, enabling simple shuttling of the carriers from HPF to cryo-FIB milling to subsequent cryo-ET imaging. After HPF, the aluminum carriers were transferred to a holder termed the Intermediate Specimen Holder (ISH) which was engineered for this application (**Figure 1c, Panel i**) to make the carrier compatible with a conventional cryo-ultramicrotome. Carriers were then trimmed, where a portion of the aluminum casing was cut away to expose ∼30 µm-thick frozen-hydrated samples (**Figure 1c, Panels ii-iii**). This cryo-trimming step facilitated cryo-FIB milling by restricting the amount of tissue required for subsequent thinning, significantly diminishing processing time and reducing risk of ice contamination. Pre-trimmed tissue was then cryo-FIB milled to obtain 150-400 nm-thick lamellae for subsequent cryo-ET imaging. A key advantage of our approach is the ability to fabricate at least 6-8 lamellae on a single carrier, each with ample surface area (*e.g*., 15 μm × 15 μm, **Figure 1e**). This provides many potential regions of interest for imaging and significantly increases the throughput of our workflow. Further, we achieved a >85% success rate, producing 130 usable tilt series from only 9 imaging sessions. Sample trimming also helps overcome a key bottleneck in cryo-ET data acquisition of cryo-FIB fabricated lamellae – a restricted tilt range^44^. Based on the geometry of the trimmed carrier at the maximal tilt angle relative to the electron beam, we calculated that a minimum distance of 80 μm (**Figure 1f-g**, orange lines) from the carrier edge is required to achieve the full tilt range of ±60°, maximizing the quality and isotropic resolution of 3D tomographic reconstructions. With these many advantages, we have been able to more easily generate cryo-tomographic data detailing biologically important subcellular structures from brain regions relevant to both health and disease.

In summary, the SISTM approach is less technically demanding and has a substantially higher success rate for generating lamellae within tissue compared to other approaches. For example, cryo-lift-out has a generally low success rate and high likelihood of ice contamination^45–47^. On the other hand, the recently developed waffle method can improve success rates for generating usable lamellae^48^. However, the waffle method’s tilt angle range is constrained, thereby limiting cryo-ET data acquisition^48^. Collectively, our SISTM method addresses the key technical bottlenecks of earlier approaches, creating a more accessible, robust workflow for *in situ* cryo-ET of tissues.

### Detailed 3D nanoarchitecture of mouse brain neuropil via SISTM

We first applied the SISTM workflow to *in situ* cryo-ET imaging of wild-type (WT) mouse brain cortical tissue. The SISTM approach offered sufficient contrast and spatial resolution to reveal distinct neuronal compartments and their organelles within brain neuropil (**Figure 2**). For example, within axonal boutons, abundant synaptic vesicles (SVs) were visualized (**Figure 2a-b**). The average diameter of these vesicles was 41.8 ± 0.7 nm, consistent with canonical SV size^49,50^. Mitochondria with extensive cristae as well as distinct outer and inner membranes were also readily identified (**Figure 2b**). Moreover, we resolved a multivesicular body (MVB) (**Figure 2c**), which contained internal vesicles with a mean diameter of 49.2 ± 1.6 nm, in line with previously published values^51,52^. Dendritic spines were also visualized, often confirmed by the presence of a spine apparatus (SA; **Figure 2d**). Spine apparatuses were characterized by multiple cisterns of presumptive smooth endoplasmic reticulum (SER) intercalated with electron-dense inner plates^53^. Additionally, Type 1 glutamatergic synapses were identified in the mouse cortex (**Figure 2e**). Type 1 synapses are distinguished by 1) a presynaptic axonal bouton filled with SVs that is 2) separated by a synaptic cleft from 3) the postsynaptic structure which contains an electron-dense postsynaptic density (PSD)^54^. The PSD is comprised of neurotransmitter receptors, cytoskeletal elements, and signal transduction proteins required for neurotransmission^55^. SISTM therefore provides a new and more accessible means to study the fundamental unit of neural communication, the synapse, under near-native conditions and in 3D.

**Figure 2.**
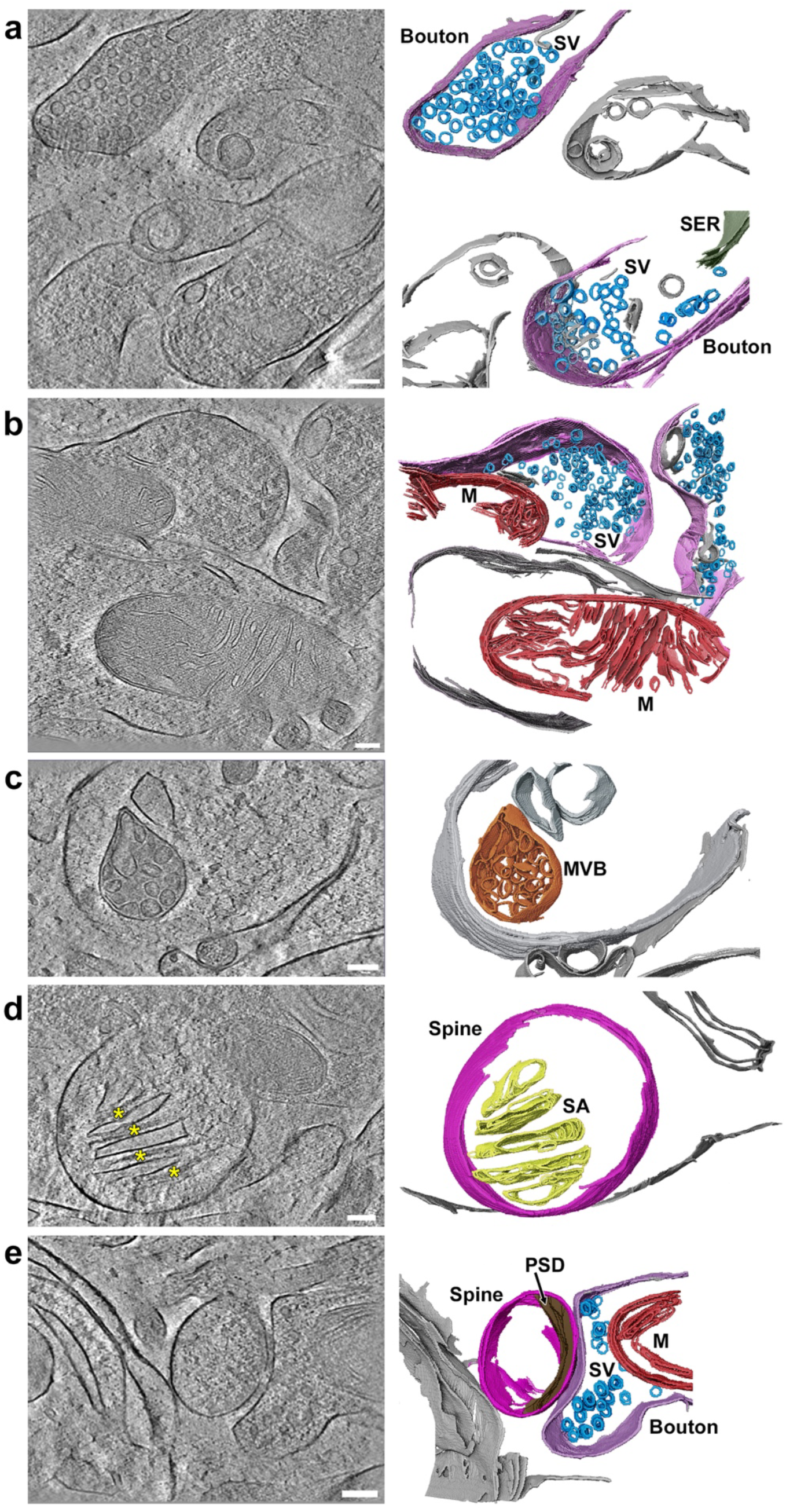
*In situ* cryo-ET of mouse cerebral cortex. Left panels show representative 2D slices from cryo-tomograms, and right panels are the corresponding segmentation of key subcellular structures. **(a)** Axonal boutons (lavender) containing clusters of synaptic vesicles (SVs, blue). The bottom bouton also contains a putative SER cisterna (olive green) alongside the SVs. **(b)** Two mitochondria (M, red) exhibit characteristic double membranes enclosing the matrix and cristae. **(c)** Multivesicular body (MVB, orange) located in a neuronal compartment. **(d)** Spine apparatus (SA, yellow) in a dendric spine (fuchsia). Characteristic electron-dense plates separating the SA cisternae are present (yellow asterisks) **(e)** Type 1 synapse. An axonal bouton (lavender), containing SVs (blue) and a mitochondrion (M, red), forms a synapse onto a dendritic spine (fuchsia) containing the postsynaptic density (PSD, brown). Scale bars = 100 nm.

### Neuronal organellar abnormalities in a Huntington’s disease mouse model

We next applied our SISTM workflow to assess neurodegenerative pathology *in situ* via cryo-ET of the R6/2 mouse model of Huntington’s disease (HD)^56^. Imaging of R6/2 mouse caudate-putamen (CPu) revealed intracellular morphological abnormalities compared to WT mice (**Figure 3a-b**). Though mitochondria from WT mice showed extensive, well-organized and thin cristae (**Figure 3a**), mitochondria from R6/2 mice exhibited dilated cristae (**Figure 3b**). We quantified cristae width and inner/outer mitochondrial distance, core mitochondrial ultrastructural features. The average cristae width in R6/2 mitochondria was significantly greater compared to WT (**Figure 3c**). On the other hand, there were no significant differences in the distance between the outer and inner mitochondrial membranes in R6/2 versus WT mice (**Figure 3d**). Our structural findings are consistent with earlier studies implicating mitochondrial dysfunction as a key feature of HD^57^, and more recent work demonstrating HD-induced alterations in cristae architecture^41,58^. Indeed, we previously demonstrated that synaptic mitochondria are specifically impacted early in R6/2 mice. The mechanism for this mitochondrial dysfunction is linked to binding of mutant huntingtin directly to mitochondrial TIM23, thus playing an early and direct role in HD pathology^59^.

**Figure 3.**
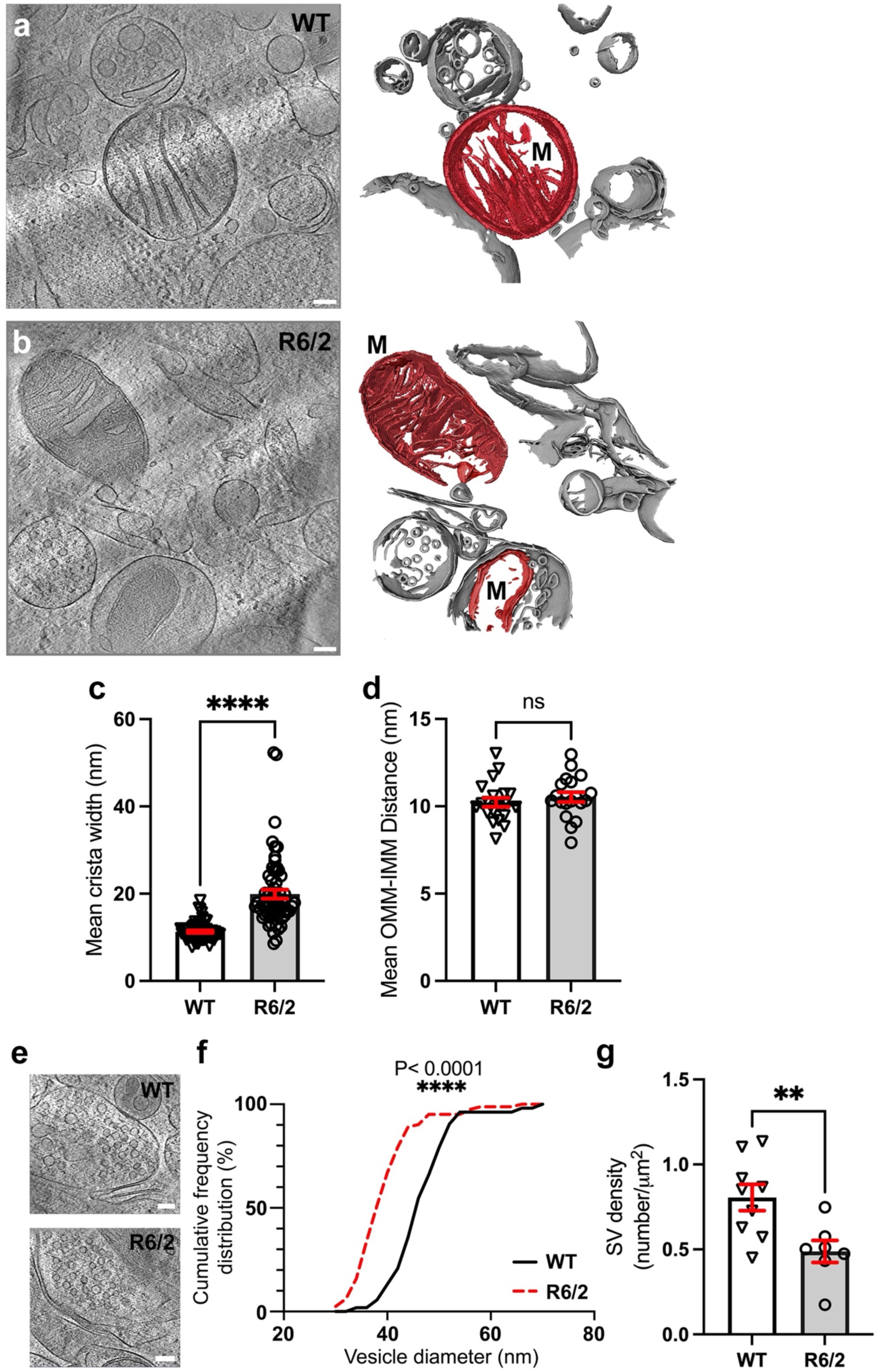
Neuronal subcellular ultrastructural abnormalities in a Huntington’s disease mouse model. **(a & b)** Caudate/putamen (CPu) from wild-type (WT) control **(a)** and Huntington’s disease model (R6/2) **(b)** mice. Representative 2D slices from cryo-tomograms (left panels) and corresponding segmentation of the subcellular features (right panels) are shown. Mitochondria in R6/2 mice exhibited noticeably wider cristae. **(c)** Quantification of mitochondrial cristae width supported this observation, with significantly wider cristae in R6/2 mice relative to WT controls. **(d)** The mean distance between outer and inner mitochondrial membranes did not significantly differ between R6/2 and WT controls. **(e)** Representative 2D slices from cryo-tomograms showing SVs from WT (top) and R6/2 (bottom) mouse CPu. **(f)** Cumulative frequency distributions for SV diameter significantly differed between WT controls and R6/2 mice. **(g)** Comparison of SV density from WT and R6/2 mouse CPu revealed significantly lower SV density in boutons. Scale bars = 100 nm. Data represented as mean ± SEM. ** p<0.01, **** p<0.0001.

Given increasing evidence of synaptic dysfunction in HD^60^, we also examined presynaptic structures in R6/2 versus WT CPu (**Figure 3e**). The average SV diameter (**Figure 3f**) and density (**Figure 3g, Extended Data Figure 2**) were significantly lower in R6/2 mice compared to WT. These morphological alterations suggest a structural basis for the abnormal glutamatergic synaptic neurotransmission reported in HD patients and mouse models^61^. Mechanistically, huntingtin protein plays a direct role in maintaining SV dynamics, including docking and recycling of SVs in boutons. Accumulation of mutant huntingtin impairs these processes^62,63^, consistent with our findings. Further work will leverage the strengths of cryo-ET to elaborate upon preexisting biochemical and functional evidence of synaptic dysfunction through the direct study of huntingtin-SV interactions directly within individually resolved synaptic units in affected tissue.

### 3D nanoarchitecture within intact macaque brain

The dorsolateral prefrontal cortex (DLPFC) is a multi-modal association area that is uniquely expanded in primates and plays a central role in mediating core processes such as cognitive control^64^. DLPFC neuronal synaptic dysfunction is strongly implicated in the pathophysiology of serious mental illness, such as psychotic spectrum disorders^65,66^. We therefore investigated individual synapses within primate DLPFC tissue in the near-native state via the SISTM workflow. Neuronal viability of the sampled DLPFC tissue from a macaque monkey was first evaluated *ex vivo*. Morphological reconstructions of glutamatergic pyramidal neurons recorded *ex vivo* revealed intact and extensive dendritic and axonal arbors (**Extended Data Figure 3a**). Consistent with these well-preserved structures, cryo-ET of DLPFC from the same monkey revealed intact myelinated axons (**Extended Data Figure 4**). Recordings from the reconstructed neurons demonstrated spontaneous excitatory postsynaptic currents (sEPSCs; **Extended Data Figure 3b**) and evoked action potential firing (**Extended Data Figure 3c**). These results confirmed that glutamatergic neuron function and effective synaptic transmission were both well-preserved in the monkey DLPFC tissue sample used for cryo-ET.

*In situ* cryo-ET imaging of DLPFC tissue from the same macaque monkey revealed a tripartite synapse (**Figure 4a-b, Supplemental Movie 2**) comprised of 1) a glutamatergic bouton forming a Type 1 synapse onto 2) a dendritic spine head, surrounded by 3) an astrocytic process that formed two physical contacts on either side of the spine head (**Figure 4c, panels i-iii**). The presynaptic bouton was densely packed with SVs, including some docked at the active zone. SVs were spherical with a mean diameter of 38.47 ± 0.47 nm (range: 30-52 nm; **Extended Data Figure 5**), consistent with earlier measurements in monkey brain^67–69^. The postsynaptic spine contained a prominent PSD, and an elaborate SA distinguished by laminated cisterns separated by dense plates (**Figure 4a-b**). Quantification of the three contact sites revealed that the Type 1 synaptic cleft width (19.02 ± 0.34 nm) was more than 50% greater than astrocyte-neuron contact 1 (10.90 ± 0.24 nm) and astrocyte-neuron contact 2 (11.90 ± 0.25 nm) clefts (**Figure 4d**), also consistent with prior measurements^70,71^.

**Figure 4.**
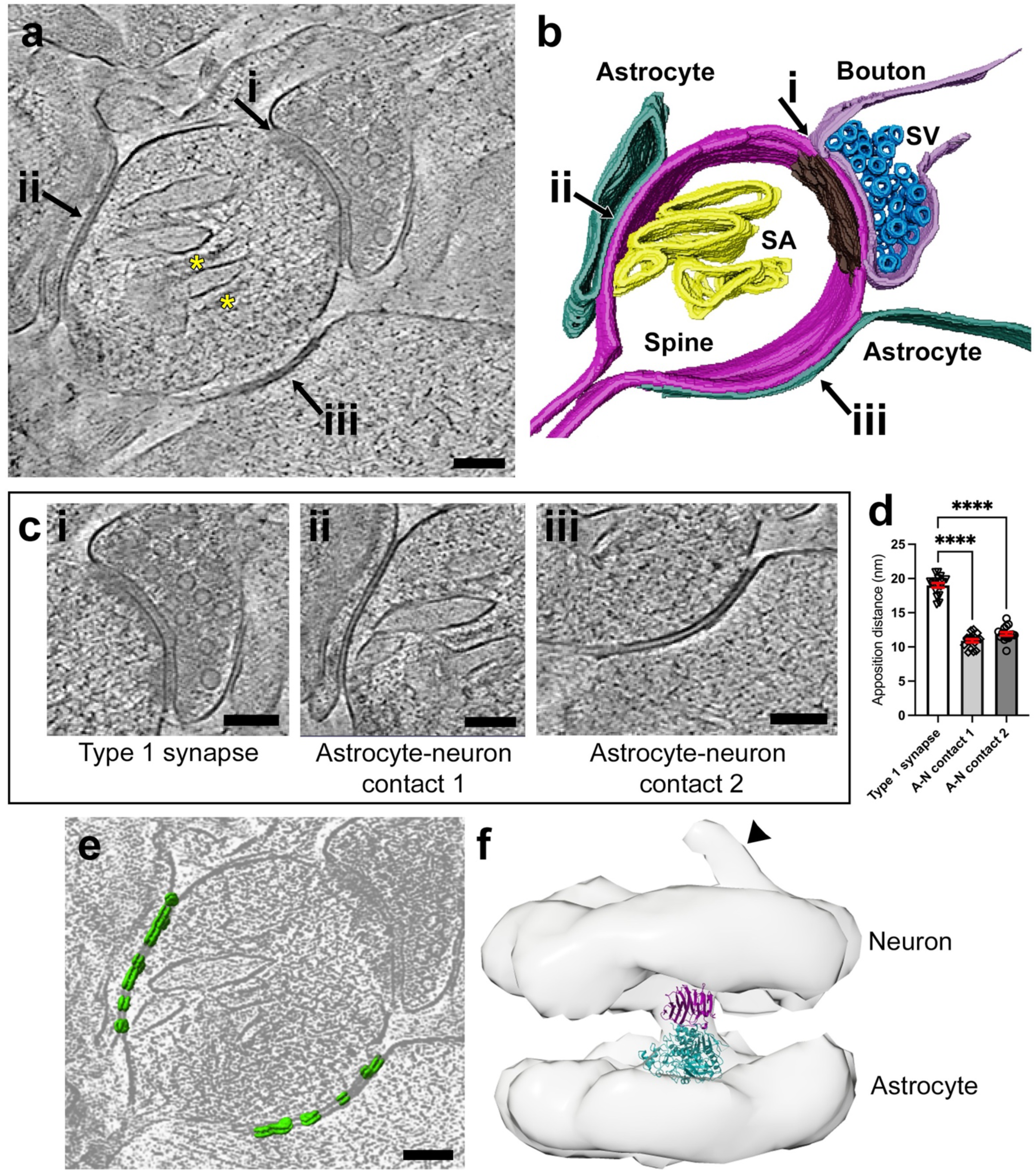
Synaptic nanoarchitecture in macaque DLPFC. **(a & b)** A representative 2D slice from cryo-tomograms **(a)** and corresponding segmentation **(b)** of a tripartite synapse. The postsynaptic dendritic spine (fuchsia) receives three contacts: i) Type 1 synapse, ii) astrocyte-neuron contact 1, and iii) astrocyte-neuron contact 2. The Type 1 synapse consists of an axonal bouton (lavender) containing SVs (blue) contacting the dendritic spine with a prominent postsynaptic density (PSD, brown). The spine also contains a spine apparatus (yellow) with interspersed electron-dense plates (yellow asterisks) separating the spine apparatus cisternae. At two distinct sites, the astrocytic process (teal) is directly apposed to the spine head and forms contacts (ii and iii). **(c)** Enlarged views of the synaptic (i) and astrocytic contacts (ii-iii). **(d)** Quantification of the apposition distances between Type 1 synapse and the two astrocyte-neuron (A-N) contacts, respectively. **(e)** Subtomogram averaged (STA) density (green) mapped into the tomogram overlaid on a cryo-ET slice, highlighting the spatial distribution along the astrocyte-neuron contacts. **(f)** STA density map of properly orientated picked particles. The map features a prominent central stalk flanked by neuron and astrocyte membranes along with a smaller density (arrowhead) protruding from the neuronal membrane into the dendritic spine head cytoplasm. Docking of the atomic model of neuroligin1-neurexin-1β complex (PDB 3BIW: neuroligin-1, teal and neurexin-1β, purple) to our STA density map revealed a good fit. Scale bars = 100 nm.

We identified electron-dense structures that were distributed throughout both astrocyte-neuron contact sites, spanning the clefts (**Figure 4e; Supplemental Movie 2**). These densities were resolved by subtomogram averaging (STA) to reveal a nanocolumn-like 3D architecture (**Figure 4e**). The initial subtomogram average showed a pseudo-2-fold symmetry, and it was difficult to assign its orientation when mapping back into the tomogram. In other words, it was not possible to ascertain which side of the averaged density belonged to the astrocyte versus the neuron. To establish the correct particle orientations, we developed a new approach that introduced an artificial fiducial on one side of the averaged density. Mapping the augmented density back into the tomogram clarified the correct orientations of the individual particles (**Extended Data Figure 6**). Correcting the orientations and refining the average again, we generated a final averaged density which revealed a slightly asymmetrical central stalk bridging the astrocyte and neuron membranes (**Figure 4f**; **Extended Data Figure 6**). We also found a smaller density emerging from the neuronal membrane into the cytoplasm of the dendritic spine head (**Figure 4f**). The Fourier Shell Correlation (FSC) curve indicated an ∼60 Å resolution at the gold-standard 0.143 threshold (**Extended Data Figure 7**). Taken together, our fiducial approach offers a new tool with which to study intrinsically asymmetric superstructures (*e.g*., different adjacent cell type-specific membranes) in 3D within their native biological systems. Just as importantly, this artificial fiducial approach offers a relatively simple, efficient method of applying directional restraints to subtomographic reconstructions.

In identifying the macromolecule(s) corresponding to the resolved 3D structure, the most logical candidates were adhesion molecules^72^. Though neurexin-neuroligin interactions are mostly studied in synaptic contacts, these molecules are also enriched at astrocyte-neuron contacts near Type 1 excitatory synapses, making them top candidates^73,74^. We therefore fitted an experimental atomic model of neuroligin-1/neurexin-1β^75^ back into the tomogram (**Figure 4f**). The atomic model fit into the density well with neurexin-1β oriented towards the neuron and neuroligin-1 oriented towards the astrocyte, consistent with prior work^73,74^ (**Figure 4f, Supplemental Movie 2**). In contrast, we also attempted to fit a LRRTM2-neurexin-1β complex^76^ into the density as a negative control given its exclusive localization to neuron-neuron synaptic contacts^77^. However, docking the atomic model into the density resulted in a considerably poorer fit that protruded into both the neuron and astrocyte membrane regions (**Extended Data Figure 8**). In addition to the central stalk, the map features a smaller density protruding from the neuronal membrane into the dendritic spine head cytoplasm (**Figure 4f arrowhead**). Presently, we do not have sufficient resolution to identify candidates for this secondary density. However, with improved resolution, further work will establish its identity.

Elucidating the identity and organization of the scaffolding underlying astrocyte-neuron contacts provides a mechanism by which astrocytes can support and maintain synaptic microenvironments. These data are also relevant to better understanding disease states in the brain. Indeed, the relationship between astrocytes and neurons at synapses was recently implicated both in normal human inter-individual differences and as a key point of convergence for multiple types of brain pathophysiology^78^. Future work will focus on examining these densities at higher resolution to definitively validate the present candidate molecules.

### Subcellular axonal bouton architecture in postmortem human brain

We next applied our SISTM workflow to examine human brain nanoarchitecture. DLPFC was obtained postmortem from a 45-year-old male who died suddenly and out-of-hospital from cardiovascular disease. This individual had no lifetime history of neuropsychiatric disorders and no neuropathologic findings. Immediately upon receipt of the brain (postmortem interval, 8.3 hours), a sample of DLPFC was excised and cryo-preserved via HPF to avoid potential artifacts associated with freeze-thawing or fixation and sectioning. Our rationale for using postmortem human brain tissue was guided by the substantial literature of rigorously conducted research studies that inform human brain structure and function across the circuit, cellular, and molecular levels^79–81^. This is coupled with a small, but growing, literature demonstrating the validity of HPF-preserved postmortem human brain tissue in cryo-ET^36,43^.

*In situ* cryo-ET imaging of human DLPFC revealed multiple components of the neuropil (**Figure 5**). Intact axonal boutons filled with SVs and containing mitochondria were readily visualized (**Figure 5a-c**). SVs were well-defined and spherical with a mean diameter of 41.38 ± 0.58 nm (range: 30-52 nm; **Figure 5d**), consistent with the measures obtained here from mouse cortex (**Figure 3f**), monkey DLPFC (**Extended Data Figure 5**), and prior work in human neocortex^82^. We identified a cluster of SVs in close apposition to the axonal bouton plasma membrane that were flanked by a putative SER cisterna (**Figure 5b**). These SVs appeared to be clathrin-coated, with visible triskelion spikes protruding from the SV membranes (**Figure 5c**). On average, spikes were 28.80 ± 1.75 nm long (range: 15-55 nm; **Figure 5e**), in line with prior measurements^83^. Clathrin-coated SVs are indicative of dynamic vesicle recycling in response to neuronal activity^83^. Presence of a cisterna-like structure alongside SVs has been previously described as either SER^84^ or an ER-derived phagophore^43^. SER/clathrin-coated vesicular microenvironments within axonal boutons likely facilitate efficient exchange of lipids between the compartments to foster vesicle recycling^85^. Since the SER serves as a major reservoir of Ca^2+^, this proximity also enables rapid local, Ca^2+^-mediated vesicle fusion^86^. Altogether, these data further support the validity of postmortem human brain tissue as an accurate representation of dynamic brain processes, as well as the ability to conduct quantitative studies via SISTM of human brain tissue in the near-native state.

**Figure 5.**
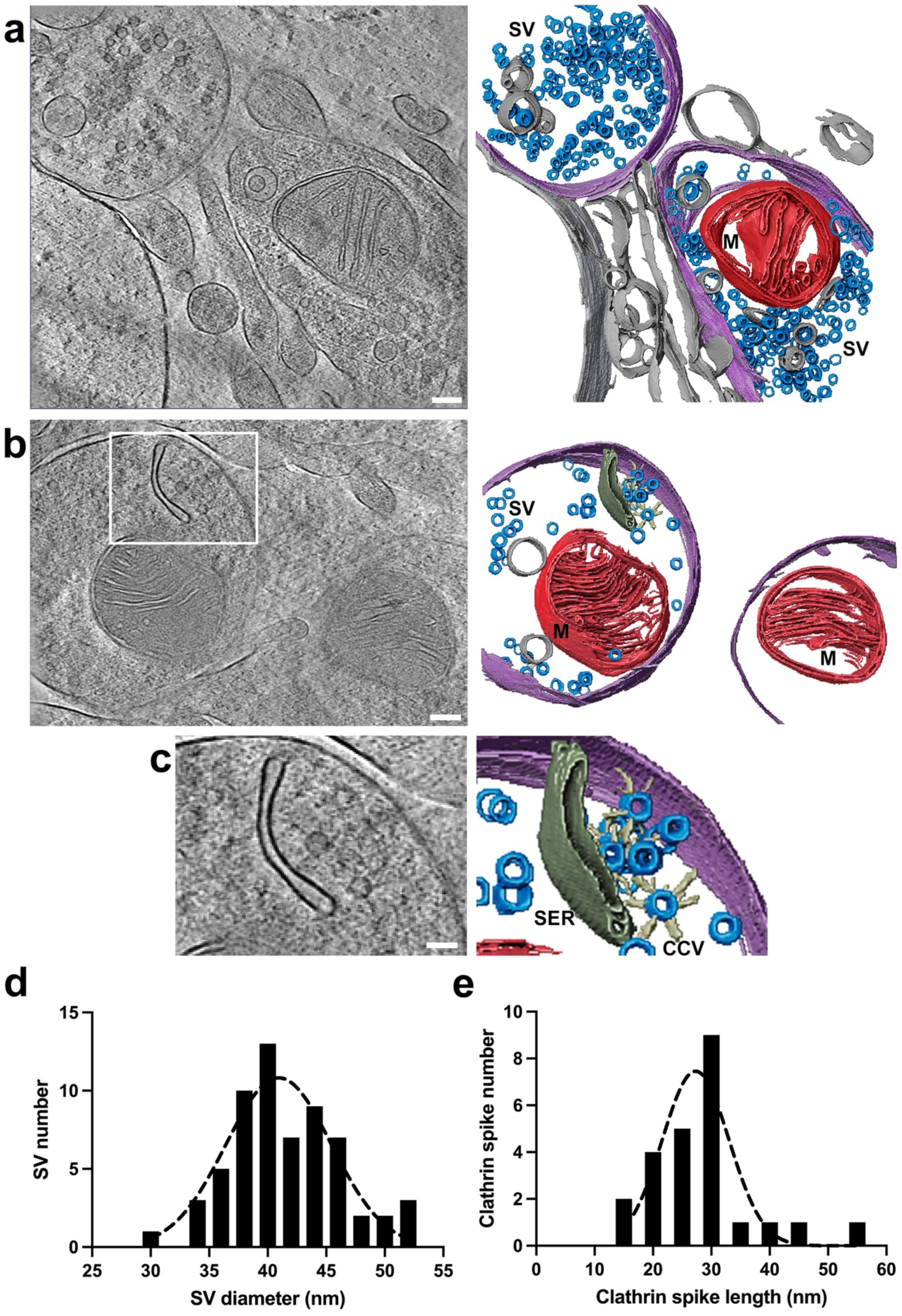
Presynaptic architecture in postmortem human DLPFC. Representative 2D slices from cryo-tomograms (left) and corresponding segmentation (right) of axonal boutons. **(a)** Two axonal boutons (lavender) filled with well-defined SVs (blue) and containing a mitochondrion (M, red). Scale bar = 100 nm. **(b)** A cluster of SVs in close apposition to the axonal bouton plasma membrane flanked by a putative SER cisterna (olive green). Scale bar = 100 nm. **(c)** Zoomed-in area of the white box indicated in (b). The SVs appear to be clathrin-coated (CCV, ivory), with putative triskelion spikes protruding from the SV membranes. Scale bar = 50 nm. **(d)** Histogram showing the distribution of mean SV diameters (N=62). **(e)** Histogram showing distribution of mean clathrin spike length (N=24).

### Conclusions

Overall, the SISTM approach significantly simplifies and increases the throughput of *in situ* cryo-ET, including in tissues as challenging as brain. This robust workflow has a low failure rate, increased throughput efficiency, and a more complete tilt range, thereby increasing the likelihood of discovering novel 3D biological data within a tissue previously challenging to image via cryogenic methods. We demonstrated the validity of this approach across different species, disease models, and brain regions. Combining SISTM with STA offers the ability to identify and visually map protein assemblies with spatial specificity in their native environments. We applied SISTM to image 3D synaptic architecture within intact cryo-preserved brain tissue from mouse, macaque, and human. In a mouse model of HD, we directly visualized the impact of HD-related pathology on presynaptic organization, finding altered 3D mitochondrial and SV structures. Within macaque DLPFC, we visualized the 3D organization of a tripartite synapse, demonstrating the nanoarchitecture of neuron-neuron and neuron-astrocyte connections. Moreover, we resolved densities within neuron-astrocyte connections, generating a novel fiducial approach for identifying putative constituents and their directionality. Finally, we show that the defining features of *in situ* synaptic architecture are conserved across species and evident in human brain tissue obtained postmortem. Overall, our approach demonstrates a promising path forward in understanding the unique subcellular nanoarchitecture of brain by combining structural data with cell biology.

## Materials and Methods

### Mice

Mice were maintained on a 12:12 hour light:dark cycle with food and water available *ad libitum*. Mice included a range of 9-29-week-old C57BL/6J (The Jackson Laboratory, Bar Harbor, ME; JAX no. 000664). The mice were anesthetized with isoflurane, transcardially perfused with phosphate-buffered saline (PBS), and then euthanized by decapitation. Brains were rapidly removed and maintained in ice-cold PBS or artificial cerebrospinal fluid (ACSF) for the duration of tissue collection. For studies using R6/2 mice, we employed male 9-week-old R6/2 (The Jackson Laboratory; JAX no. 002810) and wild-type (WT) littermates. R6/2 mice overexpressed human huntingtin exon 1 with a CAG expansion to 170 ± 10 Q as described previously^56^. The CAG repeat length was monitored every breeding cycle, and only male mice with CAG repeat lengths in this range were mated. On the experimental day, R6/2 and WT littermate mice were euthanized with carbon dioxide, transcardially perfused with cold PBS, followed by rapid removal of brains. The brains were then maintained in cold ACSF and delivered within 10 minutes of collection in ice cold state. All mouse experiments were approved by the University of Pittsburgh Institutional Animal Care and Use Committee (IACUC; protocols# 22030772, 23124171). Animals were cared for in accordance with the ARRIVE guidelines for reporting animal research^87^.

### Monkey

One male rhesus macaque aged 4 years and 9 months was maintained on a 12:12 hour light:dark cycle. The monkey was deeply anesthetized and perfused transcardially with ice-cold ACSF. The brain was removed and maintained in ice-cold ACSF for the duration of the tissue collection. All animal procedures were performed in accordance with the National Institutes of Health Guide for the Care and Use of Laboratory Animals and were approved by the University of Pittsburgh IACUC (protocol# 19024431).

### Human Subject

The brain specimen was obtained during an autopsy conducted at the Allegheny County Office of the Medical Examiner (Pittsburgh, PA) after obtaining consent from next-of-kin. An independent committee of experienced research clinicians confirmed the absence of any lifetime psychiatric or neurologic diagnoses for the decedent based on medical records, neuropathology examinations, toxicology reports, as well as structured diagnostic interviews conducted with family members of the decedent ^88,89^. The subject selected for study was a 45-year-old white male who died suddenly and out-of-hospital by natural causes from cardiovascular disease. The postmortem interval (defined as the time elapsed between death and brain tissue preservation) was 8.3 hours. Brain tissue pH was measured as 6.3, and RNA Integrity Number was determined as 8.5. These values reflect excellent tissue quality. All procedures were approved by the University of Pittsburgh’s Committee for the Oversight of Research and Clinical Training Involving Decedents and the Institutional Review Board for Biomedical Research.

### High Pressure Freezing

Brain tissue was sub-sectioned so that the region of interest was accessible for dissection. Dissected tissue was thinned and transferred directly into an aluminum carrier using a fine-tipped brush (Dynasty Brush, Glendale, NY; SC2157R, 5/0). Carriers were custom-designed for compatibility with high pressure freezing (HPF) and cryo-ultramicrotomy (Engineering Office M. Wohlwend GmbH, Sennwald, Switzerland). Carriers had a diameter of 3.0 mm, with a 0.5 mm recessed edge and a slot-shaped specimen well (1.4 mm × 0.2 mm × 0.15 mm) compatible with the Leica EMPACT2 HPF system (Leica Microsystems, Wetzlar, Germany) (see Figure 1). Immediately upon tissue transfer, carriers were loaded into a bayonet pod and torque-locked with 35.0 Ncm (Newton centimeters) of force. HPF was immediately completed at a minimum pressure of 2000 bars, and frozen samples were stored in liquid nitrogen until further use.

### Cryo-Trimming

The HPF carrier was loaded into the precooled Intermediate Specimen Holder (ISH) and transferred to the clamping chuck of the Leica UC6/FC6 cryo-ultramicrotome (Leica Microsystems) via fine forceps. Using a diamond knife (Diatome, Biel, Switzerland), a portion of the aluminum carrier was trimmed away to expose tissue. To reduce cryo-focused ion beam (cryo-FIB) milling time, the sample was further trimmed to ∼30 µm thickness (see Figure 1). After trimming, the carrier was stored under liquid nitrogen for later cryo-FIB milling.

### Cryo-FIB Milling

The pre-trimmed specimens were first clipped into FIB-autogrids by aligning the exposed tissue with the milling slot. Samples were then loaded into an Aquilos 2 cryo-FIB (Thermo Fisher Scientific/FEI, Hillsboro, OR) and milled to generate lamellae approximately 150-400 nm thickness. Detailed steps are described below:

Step 1: Once the sample was loaded, a thin platinum conductive layer was sputtered onto the sample to reduce charging. Next, an organo-platinum deposition was applied using a gas injection system (GIS) to protect the lamella during the milling process and to prevent curtaining artifacts. The sample was then rotated and tilted to face the ion beam perpendicularly. A 90° FIB milling was performed to remove a layer of tissue from the edge, preparing a smooth surface for further lamella preparation.

Step 2: Following the initial 90° cut, the exposed tissue surface was re-coated with Pt GIS. The sample was then rotated and tilted to stage positions where the FIB incidence angle ranged from 8° to 18°. At this point, the sample was prepared for progressive thinning in a stepwise manner, ensuring no curtaining artifact was introduced. The first round of thinning used 15 nA, and by the end of the process, the lamella had final dimensions of 10 µm in thickness, 200-250 µm in width, and 100 µm in depth.

Step 3: The lamella was further thinned and divided into two lamellae. Specifically, cryo-FIB milling (5 nA) of the top and bottom reduced the thickness of the lamella to 3 µm. Two lamellae were then fabricated with the final dimensions of 100 µm width and ∼60 µm depth.

Step 4: The thickness of the lamellae was reduced to 1 µm by cryo-FIB milling (0.3 nA). Six to eight 20-µm-wide lamellae were fabricated from the original two lamellae with a final depth of ∼40 µm per lamella.

Step 5: The six to eight lamellae were thinned to a 500 nm thickness by cryo-FIB milling (0.1 nA). Each lamella was ∼18 µm wide and ∼20 µm deep.

Step 6: In the final cryo-FIB milling step (50 pA), the six to eight lamellae were polished to 150-400 nm, resulting in lamellae of sufficient thinness for *in situ* cryo-ET imaging.

All cryo-FIB-milling steps were continuously monitored via scanning electron microscopy (SEM). Images were recorded with secondary electrons from the ion beam (30 kV, 10 pA) with frame integration and use of an ‘‘ETD’’ detector.

### Cryo-Electron Tomography

Samples containing lamellae were rotated 90° and loaded into an FEI Titan Krios 3Gi cryo-transmission electron microscope (TEM) equipped with a Falcon 4i direct electron detector and Selectris energy filter (Thermo Fisher Scientific/FEI). Cryo-micrographs were recorded from the cryo-FIB milled lamellae with a 3.57 Å pixel size, 300 kV electron beam, zero-loss energy filtering (20 V slit), 5-10 µm defocus. A dose symmetric tilt scheme was implemented using Tomo 5.16 software (Themo Fisher Scientific) to collect tilt series from −60° to +60° with 2° increments. The total electron dose of each tilt series was ∼100 e^-^/Å^2^.

### Tomographic Reconstruction and Annotation

Dose-fractionated video frames were imported into Warp software (v1.0.9) for frame alignment and initial contrast transfer function (CTF) estimation ^90^. Tilt series stacks were generated in Warp and imported into etomo within the IMOD software package (v4.11.25) for alignment with patch tracking. Lower quality frames were excluded with etomo before alignment. Tomograms were reconstructed via weighted back projection and simultaneous iterative reconstruction technique (SIRT)-like filtering. Tomograms were 8x-binned for segmentation with MemBrain V2 (https://github.com/teamtomo/membrain-seg)^91^ and Amira3D software (Thermo Fisher Scientific, Bordeaux, France).

### Subtomogram averaging

The macaque brain dataset was processed using RELION-5.0 (beta-4-commit-8219dd). Data were imported into the RELION-5.0 tomography pipeline, and then the raw tilt series were motion-corrected using RELION’s implementation of MotionCor2^92^. CTF was estimated using CTFFIND-4.1^93^ around the nominal defocus of 10 µm and a maximum resolution of 30 Å. Tilt series were manually inspected, and poor tilt images were removed using a Napari plug-in in the RELION-5.0 tomography pipeline^94^. Curated tilt series were automatically aligned using the IMOD wrapper for patch-tracking alignment in RELION with a patch size of 100 nm and overlap percentage of 50. The tomogram was reconstructed in RELION at a pixel size of 28.8 Å for visual inspection and particle picking.

Within the tomogram, at each astrocyte-neuron contact, electron-dense structures that spanned the contact clefts were visually identified. The coordinates of each bridging structure were obtained by drawing a line from the astrocyte membrane to the neuron membrane in the IMOD model to calculate the centroid. The Euler angles were assigned with respect to the tomogram coordinate system using a custom-written Python script, EulerMate (https://gitlab.com/ccpem). A total of 48 bridging structures from the two astrocyte-neuron contacts were picked. RELION’s Extract subtomogram job extracted the corresponding subvolumes in 32 voxel boxes at a pixel size of 14.28 Å. Using the same dataset, an initial reference was generated by running RELION’s Reconstruct particle job. The particle set was refined using Refine3D with the reference lowpass-filtered to 60 Å, without a mask, and with a particle diameter of 200 Å in C1 symmetry. The local refinements of all Euler angles were limited to 9° about priors. The tilt series was further optimized with an unbinned reference in a box size of 64 voxels without fitting per-particle motion in RELION’s Bayesian Polishing job. Particles were re-extracted from the optimized tilt series in 32 voxel boxes at a pixel size of 14.28 Å and refined as before using a spherical mask over the central bridging structure.

We next mapped the initial subtomogram averaged density back into the tomogram. The optimized particle coordinates and Euler angles were read from run_data.star from the Refine3D job. The coordinates were properly scaled to a desired pixel size and mapped into pixel coordinates. The subtomogram average was also scaled to the same pixel size. The Euler angles were converted into rotation matrices, and the subtomogram density was rotated by these matrices to obtain the correct orientation of individual particles. The rotated density was placed back into the tomogram using the pixel coordinates of the respective particles. Inspection of the particles mapped back into the tomogram revealed a total of 18 particles that did not align well with corresponding particle densities, resulting in their manual exclusion. Using the remaining 30 particles, an average density was reconstructed in a box of 32 voxels at a pixel size of 14.28 Å in RELION.

## Determining orientation of the subtomogram average with respect to the tomogram

The subtomogram average showed a pseudo-2-fold symmetry making it difficult to assign the orientation of the average with respect to the tomogram, *i.e*., determining the orientation of the averaged density relative to astrocyte and neuron sides of the respective contacts. To properly assign orientations, we introduced an artificial fiducial. The fiducial was generated by producing a cube (3 x 3 x 3 voxels with a pixel size of 28.8 Å^3^) on one side of the averaged density, which was then mapped back into the tomogram. After identifying the correct orientation using the fiducial, the particles (now without the fiducial) underwent an additional round of refinement in RELION. As before, the local refinements of all Euler angles were limited to 9° about priors. Upon convergence, the full-map was generated from the half-maps, FSC-weighted, and sharpened via EMDA software^95^.

## Docking molecular models

Model biological assemblies of candidates for docking into our subtomogram averaged density were retrieved from the Protein Data Bank (PDB). Candidates for docking included the neuroligin-1-neurexin-1β complex (PDB 3BIW)^75^, and LRRTM2-neurexin-1β complex (PDB 5Z8Y)^76^. Solvent atoms and unneeded chains were removed using PyMOL software (version 3.0; Schrödinger, L., DeLano W., 2020, http://www.pymol.org/pymol); in the case of PDB 3BIW, chains A and E were selected to form the dimer for subsequent fitting. The resultant models were docked into the density map via the “fit in map” tool in the UCSF Chimera software package^96^. The neuroligin-1-neurexin-1β complex returned a real-space correlation coefficient (RSCC) of 0.79 when positioned between the membrane regions as expected. The LRRTM2-neurexin-1β structure yielded a RSCC of 0.77. However, in contrast to the neuroligin-1-neurexin-1β complex, both ends of the LRRTM2-neurexin-1β complex clearly extend into the membrane region (Extended Data Figure 8), therefore discounting it as a plausible solution.

## Electrophysiology

### Monkey brain slice preparation

Tissue blocks containing the principal sulcus region of DLPFC area 46 from the right hemisphere were obtained from the same male rhesus macaque used for cryo-ET imaging. Tissue slices were cut at 300 μm thickness in the coronal plane starting from the rostral end of the tissue blocks using a vibrating microtome (VT1200S, Leica Microsystems) while submerged in ice-cold sucrose-artificial cerebrospinal fluid (ACSF). Immediately after cutting, the slices were transferred to an incubation chamber filled with room-temperature ACSF. Electrophysiological recordings were obtained 1-14 hours after tissue slicing was completed.

### Targeting of cells for recordings in acute brain slices

Acute brain slices were placed in recording chambers superfused at 2-3 ml/min with oxygenated ACSF at 30-32°C. To record spontaneous excitatory postsynaptic currents (sEPSCs), the GABA_A_ receptor antagonist SR95531 (10 µM, Sigma-Aldrich, St. Louis, MO) was added to the ACSF. Whole-cell recordings were obtained from pyramidal neurons identified visually by infrared differential interference contrast (IR-DIC) and epifluorescence video microscopy using Olympus or Zeiss microscopes equipped with CCD video cameras (EXi Aqua, Q-Imaging, Surrey, Canada). Cells identified as pyramidal neurons were targeted for recordings in layers 3 to 5/6 of either medial or lateral banks of the principal sulcus in DLPFC area 46.

### Electrophysiological recordings and analysis

Recording pipettes had 3-5 MΩ resistance when filled with either of the two different solutions. For voltage- and current-clamp recordings, pipettes were filled with potassium gluconate (KGluconate)-based pipette solution (composition, in mM: KGluconate 120; KCl 10; EGTA 0.2; HEPES 10; MgATP 4; NaGTP 0.3, NaPhosphocreatine 14, biocytin 0.4 %; pH 7.2-7.3, adjusted with KOH). Biocytin (0.4%) was included to fill the cells during recordings for subsequent cell visualization as described earlier ^97^.

### Voltage clamp

Recordings were obtained using Multiclamp 700A or 700B amplifiers (Axon Instruments, Union City, CA) operating in voltage clamp mode. The data were digitized at 50 kHz with Power 1401 digital-to-analog converters using Signal 5 or Signal 7 software (Cambridge Electronic Design, Cambridge, UK). Recordings were performed without employing series resistance (Rseries) compensation, while the Rseries was measured offline using in-house written Signal scripts^98^. The Rseries was monitored throughout recordings using small depolarizing or hyperpolarizing voltage steps (10 mV, 50 ms) delivered near the onset of each sweep. Sweeps were accepted for analysis only if the Rseries increased less than 20% of the initial value. Synaptic currents were recorded holding the somatic membrane potential at −80 mV. Synaptic current events were detected using NeuroMatic software^99^ with a sliding threshold search algorithm^100^ to measure: 1) event frequency (number of events detected divided by the time window analyzed); 2) the average peak amplitude; and 3) the individual sEPSC amplitudes.

### Current clamp

Recordings and data analysis were conducted with Multiclamp 700A or 700B amplifiers (Axon Instruments) operating in bridge mode with pipette capacitance neutralization^97,98^. Recordings were included in data analysis only if the resting membrane potential was ≤ −60 mV. Membrane properties were measured using families of 1 s current steps, starting from −110 pA until reaching at least 250 pA above rheobase, incrementing by 10 pA (2 repeats per current level). The procedures for measurements of membrane properties were previously described^97,98^.

## Morphological reconstruction of biocytin-filled neurons

For biocytin visualization, brain slices were resectioned at 60 μm, incubated with 1% H_2_O_2_, and immersed in blocking serum containing 0.5% Triton X-100. Sections were then incubated with the avidin–biotin–peroxidase complex (1:100; Vector Laboratories, Newark, CA), stained with the nickel-enhanced DAB chromogen, and then mounted on gelatin-coated slides, dehydrated, and cover slipped. Three-dimensional reconstructions of the dendritic arbor were performed using the Neurolucida tracing system (MBF Bioscience, Williston, VT).

## Data availability

Subtomogram averaging data will be deposited in the Protein Data Bank. Customized computer code will be deposited in Github.

## Author contributions

ZF and MM initiated the project, with ZF supervising the overall project. JN, JRG performed HPF of mouse, macaque and human brain tissue with input from JF. SAB and DC acquired fresh mouse brain tissue for the studies. JRG and DAL provided postmortem human brain tissue. JN and JRG designed and tested the carriers. TS designed and machined instrumentation specially adapted for the workflow. LGC performed cryo-trimming; JN performed cryo-FIB milling and cryo-ET imaging of brain tissue. Electrophysiology of macaque brain was performed by GGB and TM. Subtomogram averaging and model docking were performed by RW, TB, and MW. Data analysis was performed by JN, JRG, RW, TB, CMP, CH, CMH, WRS, RMF, FPM, MW, MM, and ZF. JN, JRG, RW, TB, MW and ZF wrote the manuscript with input from all the co-authors.

## Acknowledgments

This work was supported by the National Institutes of Health (R21DA052419, R21AA028800, R21AG068607 to ZF; R01NS100723 to RMF; R35GM119023 to MM, NCI cancer center support grant P30CA013330 to LGC, FPM), Commonwealth of Pennsylvania Formula Fund Award, the Pittsburgh Foundation (FPG00043-01 to ZF, RW), the Department of Defense (PR192466, PR210207 to ZF), Medical Research Council Partnership (grant No. MR/V000403/1 to TB, CP, MW), and Clear Thoughts Foundation Consortium (to RMF). The Pittsburgh Center for CryoEM (RRID:SCR_025216) used for data collection in this project was supported, in part, by the University of Pittsburgh, the School of Medicine, the Department of Structural Biology, and the National Institutes of Health (grants S10-OD-019995 and S10-OD-025009). The content is solely the responsibility of the authors and does not necessarily represent the official views of the National Institutes of Health. Postmortem human brain tissue was obtained from the NIH NeuroBioBank at the University of Pittsburgh. We are grateful to Dr. James Conway for his tremendous help and support throughout the project. We also acknowledge Dr. Donna Stolz and the Center for Biological Imaging at the University of Pittsburgh for their ongoing support throughout this work and for the assistance of Mary L. Brady in the preparation of the figures. We appreciate the input and contributions of Dr. Janet Iwasa at the University of Utah for the preparation of Supplemental Movie 2. We also thank Drs. Fei Sun, Yun Zhu, Alex Noble, and Edward Eng for discussions and technical advice.

---

**Extended Data Figure 1.**
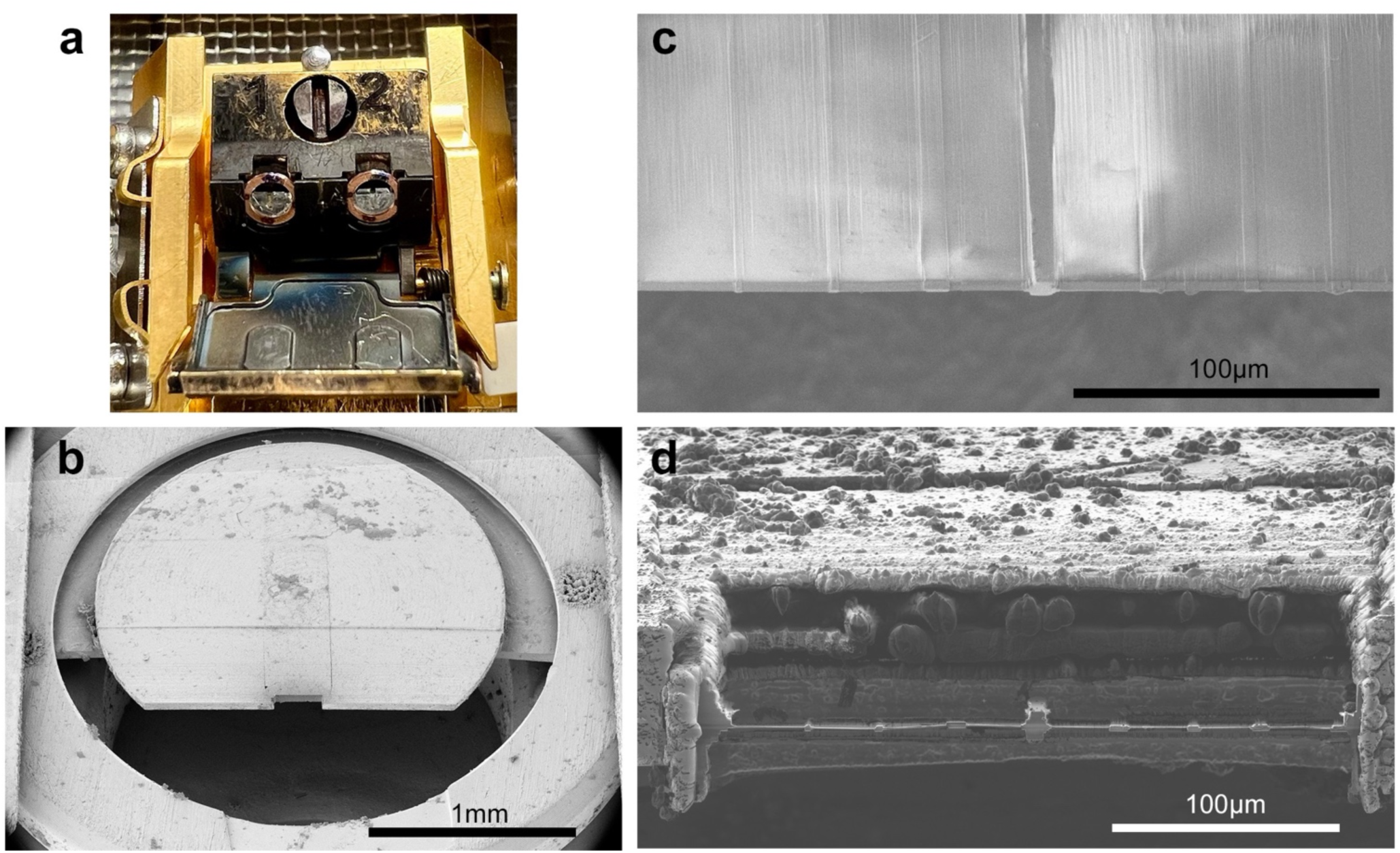
Overview of cryo-FIB milling scheme. **(a)** Two trimmed HPF carriers clipped into autogrid rings loaded into the 45° shuttle. **(b)** E-beam image of the carrier inside the dual-beam cryo-FIB chamber. Scale bar = 1 mm. **(c)** E-beam image of the manual milling scheme. Scale bar = 100 μm. **(d)** I-beam image of the 8 fabricated thin lamella. Scale bar = 100 μm.

**Extended Data Figure 2.**
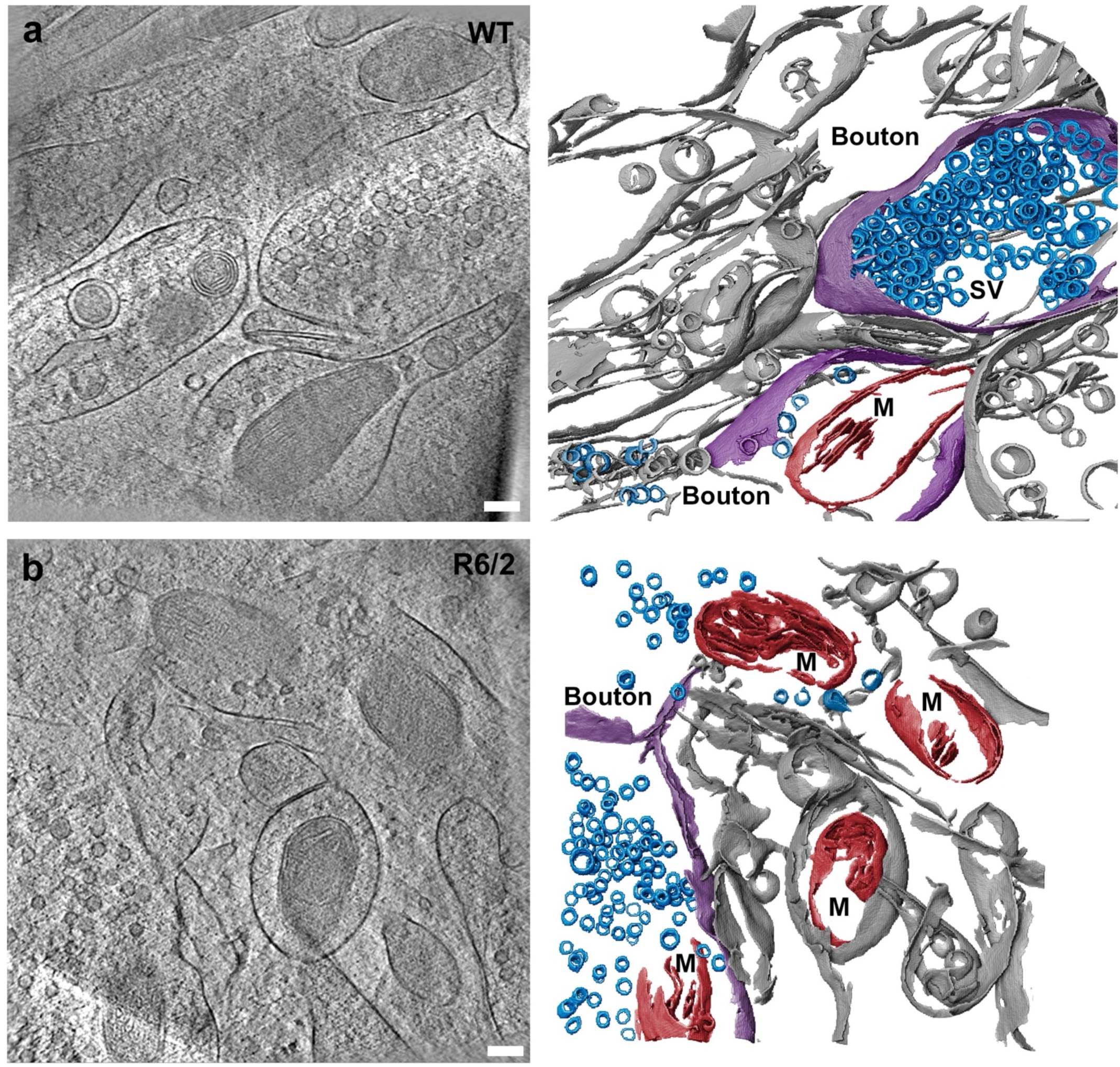
Comparation of SV distribution in WT and R6/2 mouse brain. **(a)** A representative 2D slice from cryo-tomograms and corresponding segmentation of the subcellular features of WT mouse CPu, highlighting densely packed SVs (blue) in an axonal bouton (lavender). **(b)** A representative 2D slice from cryo-tomograms and corresponding segmentation of the subcellular features of R6/2 mouse CPu, highlighting diminished SV (blue) density in an axonal bouton (lavender). Scale bars = 100 nm.

**Extended Data Figure 3.**
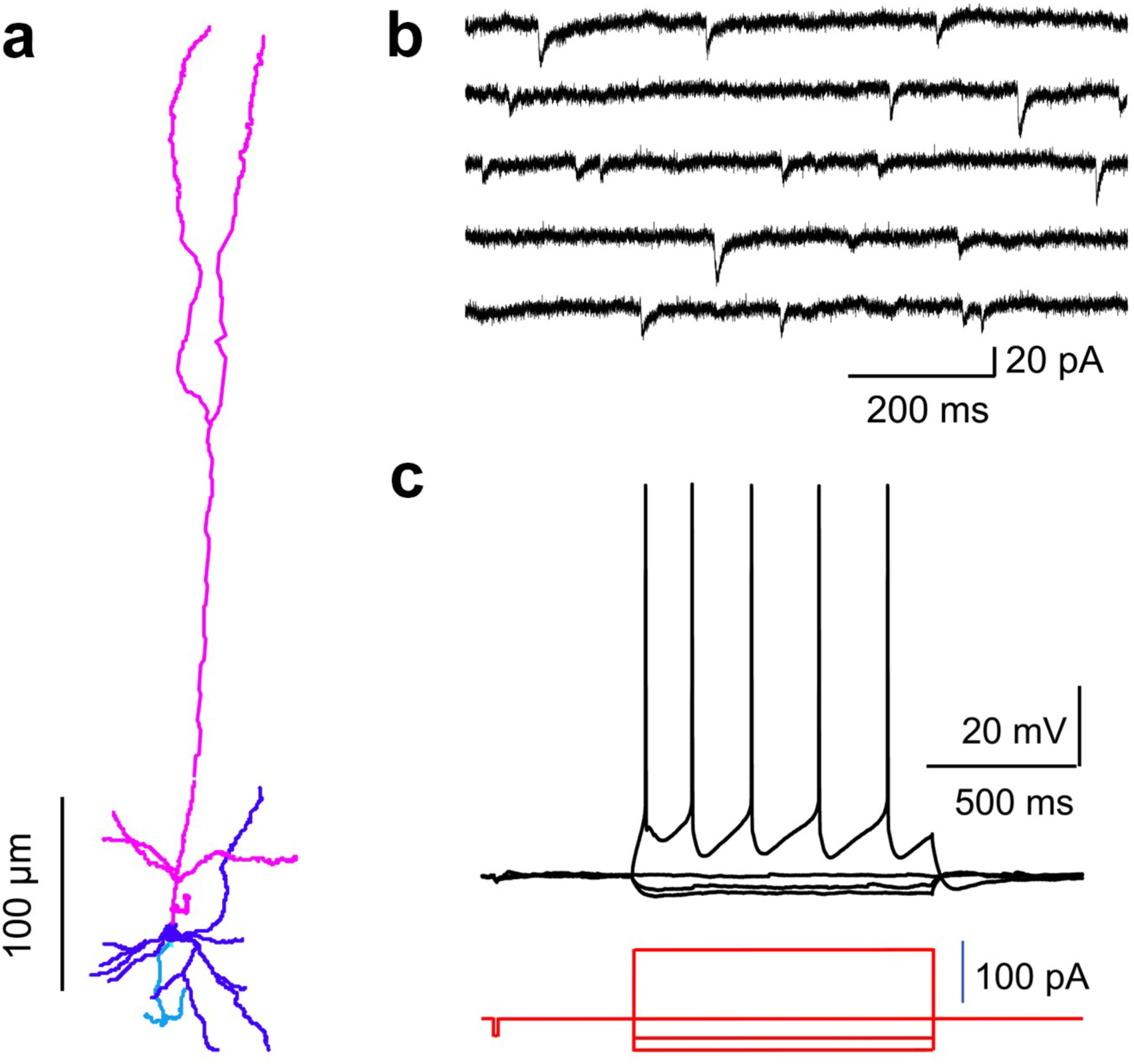
Neuronal morphological and electrophysiological properties in acute brain slices from macaque DLPFC. **(a)** Morphological reconstruction of a deep layer pyramidal neuron filled with biocytin during whole-cell patch-clamp recordings. Magenta: apical dendrite; Purple: cell body and basal dendrites; Blue: axon. **(b)** Spontaneous excitatory postsynaptic currents (sEPSCs) recorded in voltage-clamp mode from a pyramidal neuron. **(c)** Intrinsic membrane properties recorded in current-clamp mode from the pyramidal neuron in Panel a.

**Extended Data Figure 4.**
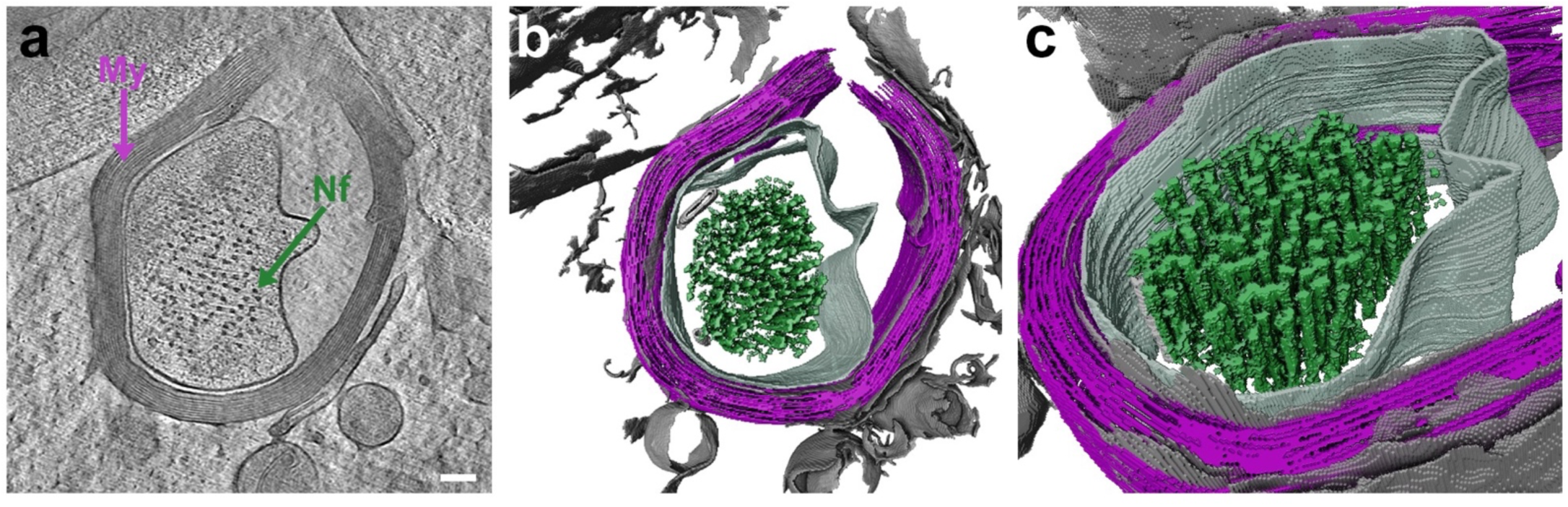
Myelinated neurons in macaque DLPFC. **(a)** A representative 2D slice from cryo-tomograms of fresh, unfixed macaque DLPFC illustrating a myelinated axon. Neurofilaments (Nf) fill the central axon which is wrapped by multiple layers of myelin (My). **(b)** Corresponding segmentation of the myelinated axon in Panel a. Neurofilaments (green) fill the central axon, which is ensheathed by myelin (purple). **(c)** Zoomed-in view of the neurofilaments and myelin from Panel b.

**Extended Data Figure 5.**
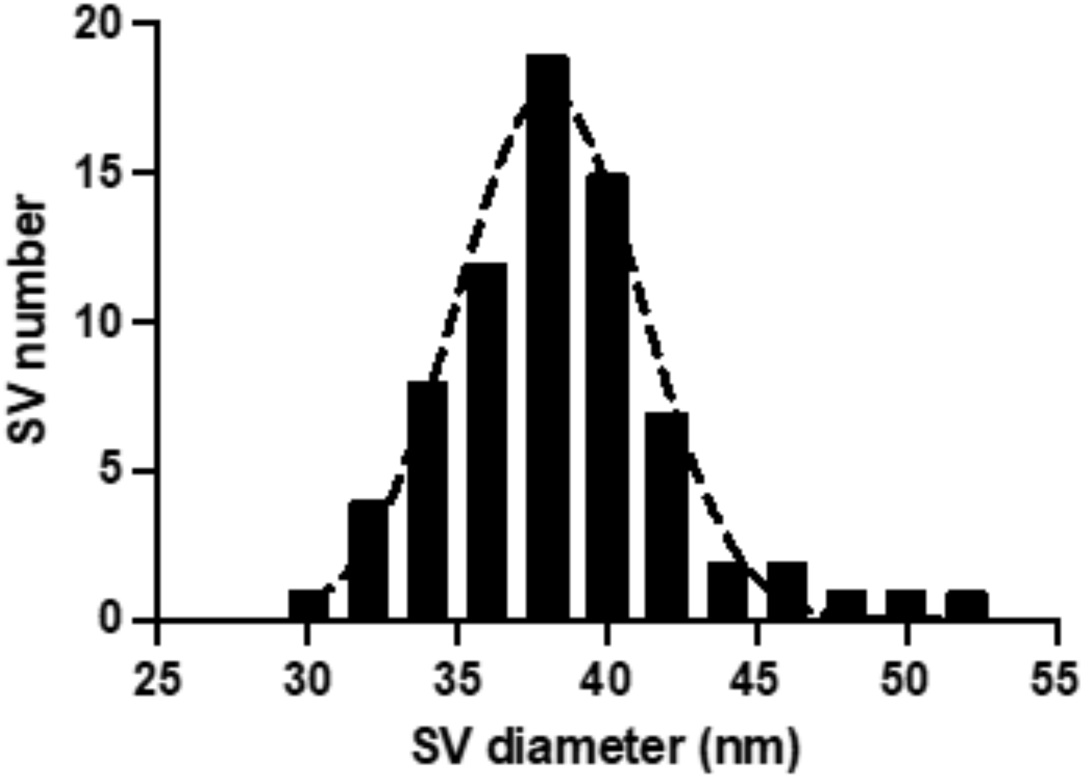
The distribution of mean axonal bouton SV diameters in macaque DLPFC. SV (N = 73) diameters were measured in *in situ* cryo-tomograms of fresh, unfixed macaque DLPFC, as represented in a histogram showing diameter distribution.

**Extended Data Figure 6.**
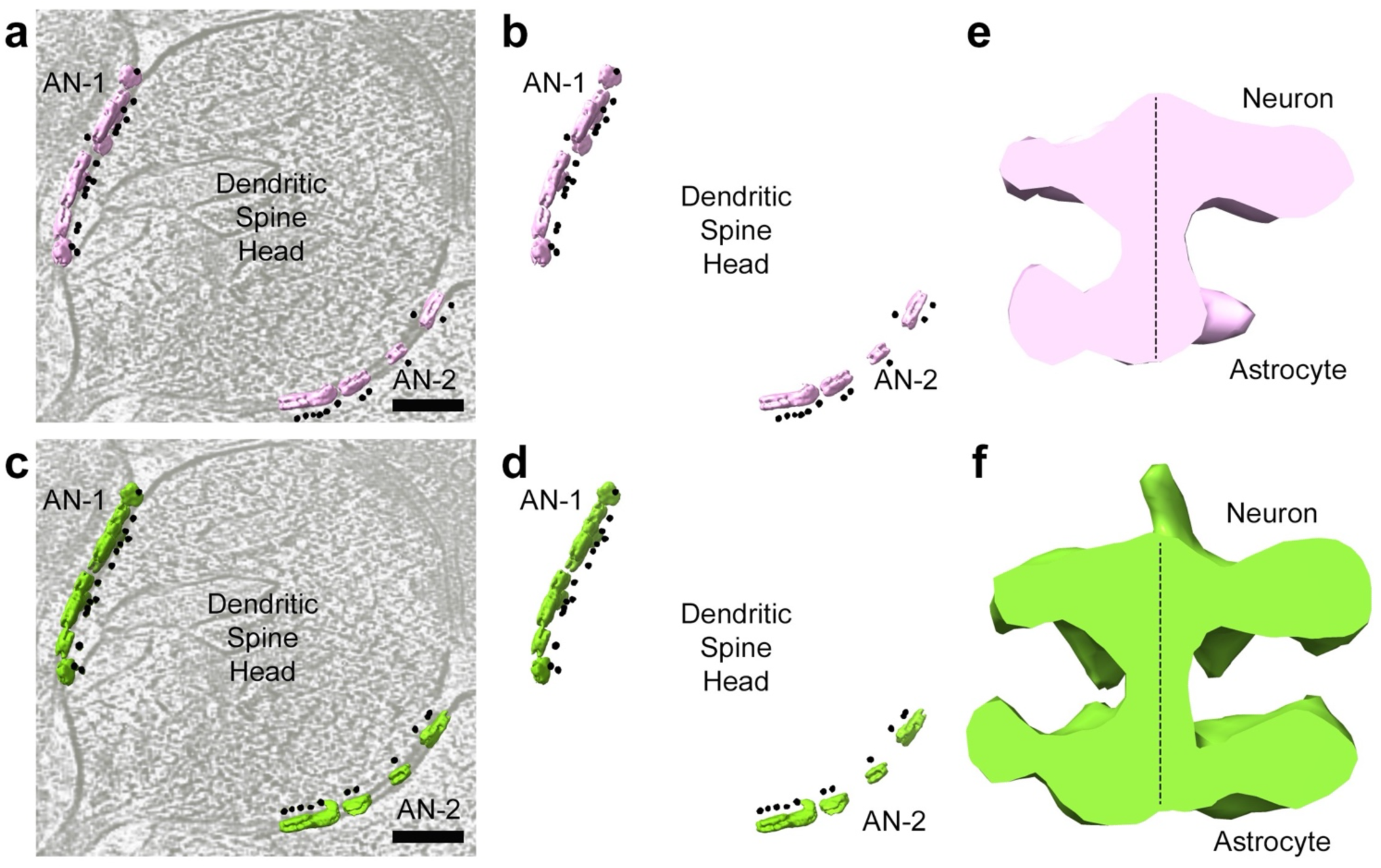
Fiducial-based orientation of the averaged density relative to neuron and astrocyte membranes. **(a & b)** The averaged density with an associated artificial fiducial is mapped back into the cryo-tomogram **(a)**, and in isolation **(b)**. Note that almost all particles associated with astrocyte-neuron contact 1 (AN-1) face the astrocyte, while the particles associated with astrocyte-neuron contact 2 (AN-2) face the spine head (also see Figure 4c). **(c & d)** The fiducial-assisted reorientation places the associated artificial fiducial uniformly on the neuron side facing the spine head, as shown mapped back into the tomogram **(c)** and in isolation **(d)**. **(e & f)** The subtomogram averaged densities corresponding to particle orientations in panels **a** and **c**, respectively. Note the slight asymmetry in the properly oriented density (green) compared to that of the original non-uniformly oriented density (pink) about the stem. The asymmetry may result from the reorientation of the particles. Scale bars = 100 nm.

**Extended Data Figure 7.**
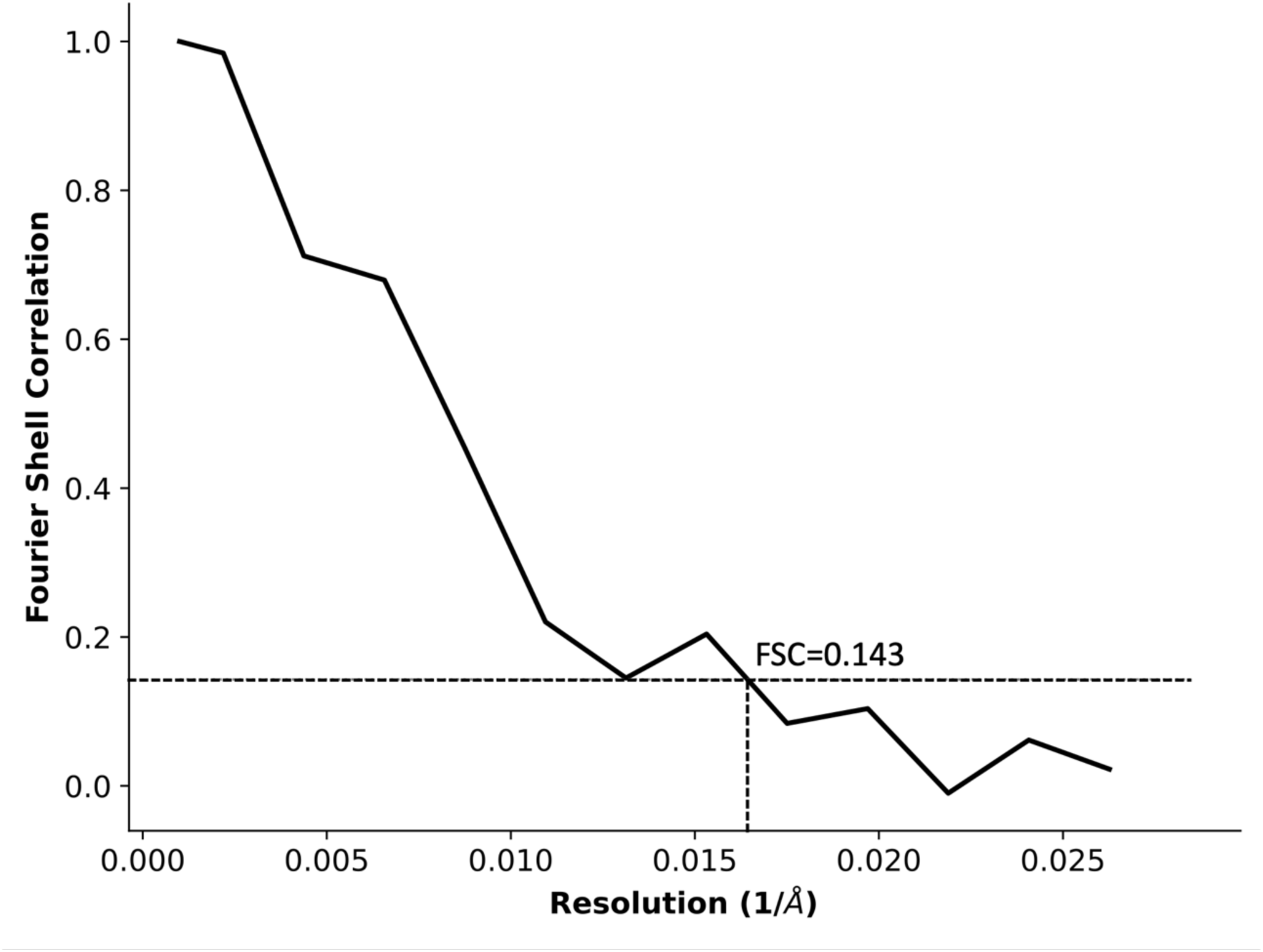
Estimation of resolution of the subtomogram averaged density. Fourier shell correlation (FSC) curve of subtomogram averaged densities shown in Figure 4e-f. The gold-standard 0.143 cut-off provided an estimate of ∼60 Å resolution.

**Extended Figure 8.**
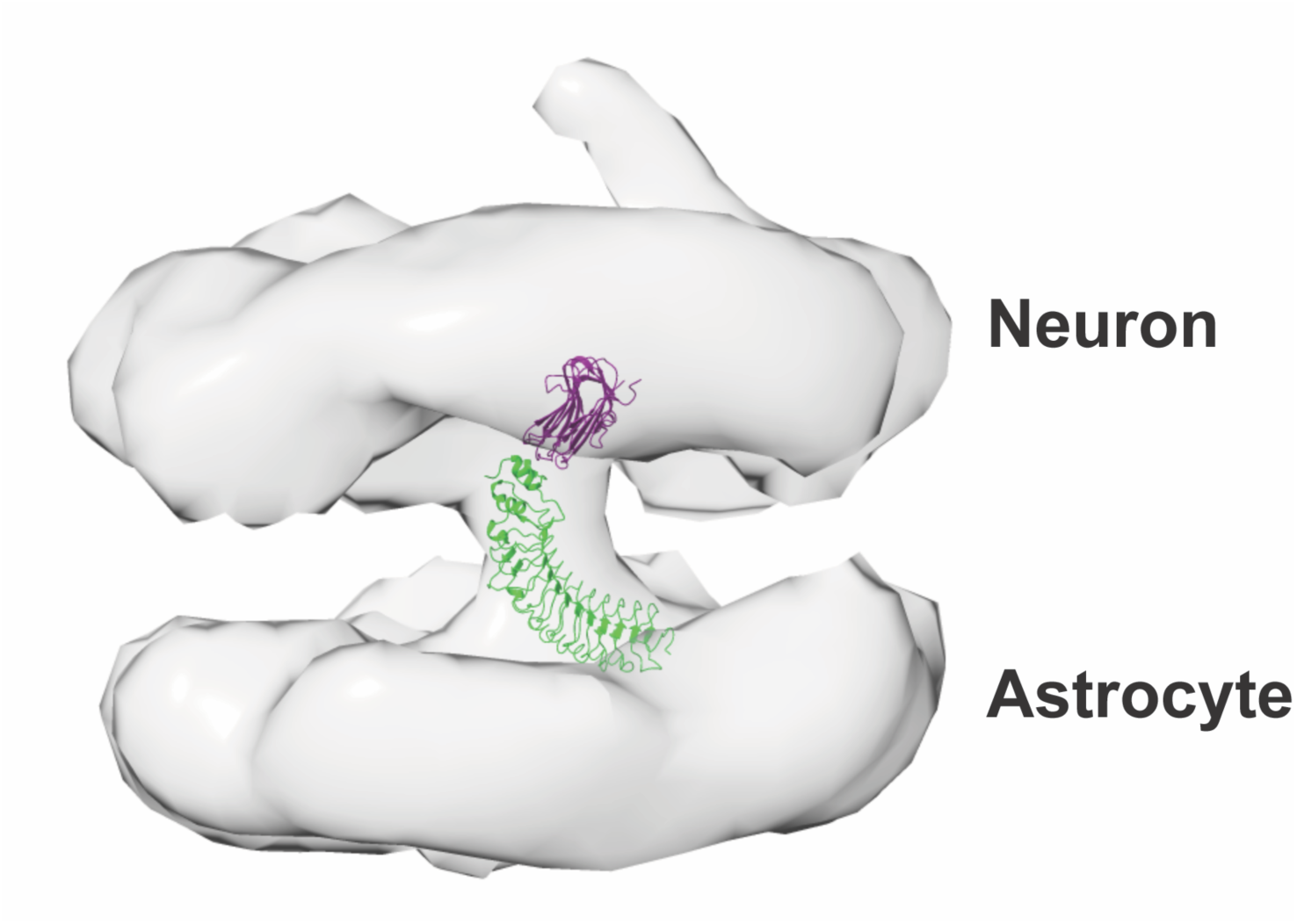
Docking the atomic model of LRRTM2/neurexin-1β to the subtomogram averaged density map. Docking the cell adhesion complex LRRTM2 (green)/neurexin-1β (purple) (PDB 5Z8Y) into the subtomogram averaged density map.

## Supplemental Movies

Supplemental Movie 1. *In situ* cryo-ET of mouse cerebral cortex.

Supplemental Movie 2. *In situ* cryo-ET of a tripartite synapse in macaque dorsolateral prefrontal cortex.

Supplemental Movie 3. *In situ* cryo-ET of presynaptic architecture in postmortem human dorsolateral prefrontal cortex.

## References

1 Wu, M. & Lander, G. C. Present and Emerging Methodologies in Cryo-EM Single-Particle Analysis. Biophys J 119, 1281–1289 (2020). 10.1016/j.bpj.2020.08.027

2 Lyumkis, D. Challenges and opportunities in cryo-EM single-particle analysis. J Biol Chem 294, 5181–5197 (2019). 10.1074/jbc.REV118.005602

3 Oikonomou, C. M. & Jensen, G. J. Cellular Electron Cryotomography: Toward Structural Biology In Situ. Annu Rev Biochem (2017). 10.1146/annurev-biochem-061516-044741

4 Berger, C. et al. Cryo-electron tomography on focused ion beam lamellae transforms structural cell biology. Nat Methods 20, 499–511 (2023). 10.1038/s41592-023-01783-5

5 Asarnow, D. et al. Recent advances in infectious disease research using cryo-electron tomography. Front Mol Biosci 10, 1296941 (2023). 10.3389/fmolb.2023.1296941

6 Young, L. N. & Villa, E. Bringing Structure to Cell Biology with Cryo-Electron Tomography. Annu Rev Biophys 52, 573–595 (2023). 10.1146/annurev-biophys-111622-091327

7 Kixmoeller, K., Creekmore, B. C., Lee, E. B. & Chang, Y. W. Bridging structural biology and clinical research through in-tissue cryo-electron tomography. The EMBO journal 43, 4810–4813 (2024). 10.1038/s44318-024-00216-z

8 Lopes-Ramos, C. M. et al. Regulatory network changes between cell lines and their tissues of origin. BMC Genomics 18, 723 (2017). 10.1186/s12864-017-4111-x

9 Campiglio, C. E. et al. Influence of Culture Substrates on Morphology and Function of Pulmonary Alveolar Cells In Vitro. Biomolecules 11 (2021). 10.3390/biom11050675

10 Hakkinen, K. M., Harunaga, J. S., Doyle, A. D. & Yamada, K. M. Direct comparisons of the morphology, migration, cell adhesions, and actin cytoskeleton of fibroblasts in four different three-dimensional extracellular matrices. Tissue Eng Part A 17, 713–724 (2011). 10.1089/ten.TEA.2010.0273

11 Keil, K. P., Sethi, S., Wilson, M. D., Chen, H. & Lein, P. J. In vivo and in vitro sex differences in the dendritic morphology of developing murine hippocampal and cortical neurons. Sci Rep 7, 8486 (2017). 10.1038/s41598-017-08459-z

12 Dauth, S. et al. Neurons derived from different brain regions are inherently different in vitro: a novel multiregional brain-on-a-chip. Journal of neurophysiology 117, 1320–1341 (2017). 10.1152/jn.00575.2016

13 Ho, V. M., Lee, J. A. & Martin, K. C. The cell biology of synaptic plasticity. Science 334, 623–628 (2011). 10.1126/science.1209236

14 Kulik, Y. D., Watson, D. J., Cao, G., Kuwajima, M. & Harris, K. M. Structural plasticity of dendritic secretory compartments during LTP-induced synaptogenesis. eLife 8 (2019). 10.7554/eLife.46356

15 Bailey, C. H., Kandel, E. R. & Harris, K. M. Structural Components of Synaptic Plasticity and Memory Consolidation. Cold Spring Harb Perspect Biol 7, a021758 (2015). 10.1101/cshperspect.a021758

16 Megías, M., Emri, Z., Freund, T. F. & Gulyás, A. I. Total number and distribution of inhibitory and excitatory synapses on hippocampal CA1 pyramidal cells. Neuroscience 102, 527–540 (2001). 10.1016/s0306-4522(00)00496-6

17 DeFelipe, J. et al. Neocortical circuits: evolutionary aspects and specificity versus non-specificity of synaptic connections. Remarks, main conclusions and general comments and discussion. J Neurocytol 31, 387–416 (2002). 10.1023/a:1024142513991

18 DeFelipe, J., Alonso-Nanclares, L. & Arellano, J. I. Microstructure of the neocortex: comparative aspects. J Neurocytol 31, 299–316 (2002). 10.1023/a:1024130211265

19 Kandel, E. R. The molecular biology of memory storage: a dialogue between genes and synapses. Science 294, 1030–1038 (2001). 10.1126/science.1067020

20 Liu, Y. et al. Interactions of glial cells with neuronal synapses, from astrocytes to microglia and oligodendrocyte lineage cells. Glia 71, 1383–1401 (2023). 10.1002/glia.24343

21 Rangel-Gomez, M. et al. Neuron-Glial Interactions: Implications for Plasticity, Behavior, and Cognition. J Neurosci 44 (2024). 10.1523/jneurosci.1231-24.2024

22 Glausier, J. R. & Lewis, D. A. Dendritic spine pathology in schizophrenia. Neuroscience 251, 90–107 (2013). 10.1016/j.neuroscience.2012.04.044

23 Glausier, J. R. & Lewis, D. A. Mapping pathologic circuitry in schizophrenia. Handb Clin Neurol 150, 389–417 (2018). 10.1016/b978-0-444-63639-3.00025-6

24 van Spronsen, M. & Hoogenraad, C. C. Synapse pathology in psychiatric and neurologic disease. Curr Neurol Neurosci Rep 10, 207–214 (2010). 10.1007/s11910-010-0104-8

25 Marsden, W. N. Synaptic plasticity in depression: molecular, cellular and functional correlates. Progress in neuro-psychopharmacology & biological psychiatry 43, 168–184 (2013). 10.1016/j.pnpbp.2012.12.012

26 Marín, O. Interneuron dysfunction in psychiatric disorders. Nat Rev Neurosci 13, 107–120 (2012). 10.1038/nrn3155

27. Dhuriya, Y. K. & Sharma, D. Neuronal Plasticity: Neuronal Organization is Associated with Neurological Disorders. Journal of molecular neuroscience : MN 70, 1684-1701 (2020). 10.1007/s12031-020-01555-2

28 Jackson-Lewis, V., Jakowec, M., Burke, R. E. & Przedborski, S. Time course and morphology of dopaminergic neuronal death caused by the neurotoxin 1-methyl-4-phenyl-1,2,3,6-tetrahydropyridine. Neurodegeneration : a journal for neurodegenerative disorders, neuroprotection, and neuroregeneration 4, 257–269 (1995).

29 Marko, M., Hsieh, C., Moberlychan, W., Mannella, C. A. & Frank, J. Focused ion beam milling of vitreous water: prospects for an alternative to cryo-ultramicrotomy of frozen-hydrated biological samples. Journal of microscopy 222, 42–47 (2006). 10.1111/j.1365-2818.2006.01567.x

30 Marko, M., Hsieh, C., Schalek, R., Frank, J. & Mannella, C. Focused-ion-beam thinning of frozen-hydrated biological specimens for cryo-electron microscopy. Nat Methods 4, 215–217 (2007). 10.1038/nmeth1014

31 Strunk, K. M., Wang, K., Ke, D., Gray, J. L. & Zhang, P. Thinning of large mammalian cells for cryo-TEM characterization by cryo-FIB milling. Journal of microscopy 247, 220–227 (2012). 10.1111/j.1365-2818.2012.03635.x

32 Rigort, A. et al. Focused ion beam micromachining of eukaryotic cells for cryoelectron tomography. Proc Natl Acad Sci U S A 109, 4449–4454 (2012). 10.1073/pnas.1201333109

33 Zhang, J. et al. VHUT-cryo-FIB, a method to fabricate frozen hydrated lamellae from tissue specimens for in situ cryo-electron tomography. J Struct Biol 213, 107763 (2021). 10.1016/j.jsb.2021.107763

34 Hsieh, C., Schmelzer, T., Kishchenko, G., Wagenknecht, T. & Marko, M. Practical workflow for cryo focused-ion-beam milling of tissues and cells for cryo-TEM tomography. J Struct Biol 185, 32–41 (2014). 10.1016/j.jsb.2013.10.019

35 He, J. et al. Cryo-FIB specimen preparation for use in a cartridge-type cryo-TEM. J Struct Biol 199, 114–119 (2017). 10.1016/j.jsb.2017.05.011

36 Gilbert, M. A. G. et al. CryoET of β-amyloid and tau within postmortem Alzheimer’s disease brain. Nature 631, 913–919 (2024). 10.1038/s41586-024-07680-x

37 Matsui, A. et al. Cryo-electron tomographic investigation of native hippocampal glutamatergic synapses. eLife 13 (2024). 10.7554/eLife.98458

38 Xu, J. et al. In situ structural insights into the excitation-contraction coupling mechanism of skeletal muscle. Sci Adv 10, eadl1126 (2024). 10.1126/sciadv.adl1126

39 Schaffer, M. et al. A cryo-FIB lift-out technique enables molecular-resolution cryo-ET within native Caenorhabditis elegans tissue. Nat Methods 16, 757–762 (2019). 10.1038/s41592-019-0497-5

40 Kuba, J. et al. Advanced cryo-tomography workflow developments - correlative microscopy, milling automation and cryo-lift-out. Journal of microscopy 281, 112–124 (2021). 10.1111/jmi.12939

41 Wu, G. H. et al. CryoET reveals organelle phenotypes in huntington disease patient iPSC-derived and mouse primary neurons. Nature communications 14, 692 (2023). 10.1038/s41467-023-36096-w

42 Martinez-Sanchez, A. et al. Trans-synaptic assemblies link synaptic vesicles and neuroreceptors. Sci Adv 7 (2021). 10.1126/sciadv.abe6204

43 Creekmore, B. C., Kixmoeller, K., Black, B. E., Lee, E. B. & Chang, Y. W. Ultrastructure of human brain tissue vitrified from autopsy revealed by cryo-ET with cryo-plasma FIB milling. Nature communications 15, 2660 (2024). 10.1038/s41467-024-47066-1

44 Noble, A. J. & de Marco, A. Cryo-focused ion beam for in situ structural biology: State of the art, challenges, and perspectives. Current Opinion in Structural Biology 87, 102864 (2024). 10.1016/j.sbi.2024.102864

45 Schiøtz, O. H. et al. Serial Lift-Out: sampling the molecular anatomy of whole organisms. Nat Methods 21, 1684–1692 (2024). 10.1038/s41592-023-02113-5

46 Parmenter, C. D., Fay, M. W., Hartfield, C. & Eltaher, H. M. Making the practically impossible “Merely difficult”--Cryogenic FIB lift-out for “Damage free” soft matter imaging. Microsc Res Tech 79, 298–303 (2016). 10.1002/jemt.22630

47 Parmenter, C. D. & Nizamudeen, Z. A. Cryo-FIB-lift-out: practically impossible to practical reality. Journal of microscopy 281, 157–174 (2021). 10.1111/jmi.12953

48 Kelley, K. et al. Waffle Method: A general and flexible approach for improving throughput in FIB-milling. Nature Communications 13, 1857 (2022). 10.1038/s41467-022-29501-3

49 Südhof, T. C. in Basic Neurochemistry: Molecular, Cellular and Medical Aspects. 6th edition. (1999).

50 Zhang, B. et al. Synaptic Vesicle Size and Number Are Regulated by a Clathrin Adaptor Protein Required for Endocytosis. Neuron 21, 1465–1475 (1998). 10.1016/S0896-6273(00)80664-9

51 Von Bartheld, C. S. & Altick, A. L. Multivesicular bodies in neurons: distribution, protein content, and trafficking functions. Prog Neurobiol 93, 313–340 (2011). 10.1016/j.pneurobio.2011.01.003

52 Oikonomou, C. M. & Jensen, G. J. Cellular Electron Cryotomography: Toward Structural Biology In Situ. Annu Rev Biochem 86, 873–896 (2017). 10.1146/annurevbiochem-061516-044741

53 Falahati, H., Wu, Y., Fang, M. & De Camilli, P. Ectopic reconstitution of a spine-apparatus-like structure provides insight into mechanisms underlying its formation. Curr Biol 35, 265–276.e264 (2025). 10.1016/j.cub.2024.11.010

54 Peters, A., Palay, S. L. & deF Webster, H. *The fine structure of the nervous system: neurons and their supporting cells*. Third ed. edn, (Oxford University Press, 1991).

55 Droogers, W. J. & MacGillavry, H. D. Plasticity of postsynaptic nanostructure. Molecular and cellular neurosciences 124, 103819 (2023). 10.1016/j.mcn.2023.103819

56 Mangiarini, L. et al. Exon 1 of the HD gene with an expanded CAG repeat is sufficient to cause a progressive neurological phenotype in transgenic mice. Cell 87, 493–506 (1996). 10.1016/s0092-8674(00)81369-0

57 Quintanilla, R. A. & Johnson, G. V. Role of mitochondrial dysfunction in the pathogenesis of Huntington’s disease. Brain Res Bull 80, 242–247 (2009). 10.1016/j.brainresbull.2009.07.010

58 Martin-Solana, E. et al. Disruption of the mitochondrial network in a mouse model of Huntington’s disease visualized by in-tissue multiscale 3D electron microscopy. Acta Neuropathol Commun 12, 88 (2024). 10.1186/s40478-024-01802-2

59 Yano, H. et al. Inhibition of mitochondrial protein import by mutant huntingtin. Nat Neurosci 17, 822–831 (2014). 10.1038/nn.3721

60 Cepeda, C. & Levine, M. S. Synaptic Dysfunction in Huntington’s Disease: Lessons from Genetic Animal Models. Neuroscientist 28, 20–40 (2022). 10.1177/1073858420972662

61 Barron, J. C., Hurley, E. P. & Parsons, M. P. Huntingtin and the Synapse. Front Cell Neurosci 15, 689332 (2021). 10.3389/fncel.2021.689332

62 Li, H., Wyman, T., Yu, Z. X., Li, S. H. & Li, X. J. Abnormal association of mutant huntingtin with synaptic vesicles inhibits glutamate release. Hum Mol Genet 12, 2021–2030 (2003). 10.1093/hmg/ddg218

63 Chen, S. et al. Real-time three-dimensional tracking of single vesicles reveals abnormal motion and pools of synaptic vesicles in neurons of Huntington’s disease mice. iScience 24, 103181 (2021). 10.1016/j.isci.2021.103181

64 Lesh, T. A., Niendam, T. A., Minzenberg, M. J. & Carter, C. S. Cognitive control deficits in schizophrenia: mechanisms and meaning. Neuropsychopharmacology 36, 316–338 (2011). 10.1038/npp.2010.156

65 Vinogradov, S., Chafee, M. V., Lee, E. & Morishita, H. Psychosis spectrum illnesses as disorders of prefrontal critical period plasticity. Neuropsychopharmacology 48, 168–185 (2023). 10.1038/s41386-022-01451-w

66 Smucny, J., Dienel, S. J., Lewis, D. A. & Carter, C. S. Mechanisms underlying dorsolateral prefrontal cortex contributions to cognitive dysfunction in schizophrenia. Neuropsychopharmacology 47, 292–308 (2022). 10.1038/s41386-021-01089-0

67 Knyihar-Csillik, E., Csillik, B. & Rakic, P. Ultrastructure of normal and degenerating glomerular terminals of dorsal root axons in the substantia gelatinosa of the rhesus monkey. J Comp Neurol 210, 357–375 (1982). 10.1002/cne.902100404

68 Cooper, M. H. & Beal, J. A. The neurons and the synaptic endings in the primate basilar pontine gray. J Comp Neurol 180, 17–41 (1978). 10.1002/cne.901800103

69 DiFiglia, M., Aronin, N. & Leeman, S. E. Light microscopic and ultrastructural localization of immunoreactive substance P in the dorsal horn of monkey spinal cord. Neuroscience 7, 1127–1139 (1982). 10.1016/0306-4522(82)91120-4

70 Torres-Ceja, B. & Olsen, M. L. A closer look at astrocyte morphology: Development, heterogeneity, and plasticity at astrocyte leaflets. Curr Opin Neurobiol 74, 102550 (2022). 10.1016/j.conb.2022.102550

71 Medvedev, N. et al. Glia selectively approach synapses on thin dendritic spines. Philosophical transactions of the Royal Society of London. Series B, Biological sciences 369, 20140047 (2014). 10.1098/rstb.2014.0047

72 Ngoc, K. H., Jeon, Y., Ko, J. & Um, J. W. Multifarious astrocyte-neuron dialog in shaping neural circuit architecture. Trends Cell Biol 35, 74–87 (2025). 10.1016/j.tcb.2024.05.002

73 Saint-Martin, M. & Goda, Y. Astrocyte-synapse interactions and cell adhesion molecules. The FEBS journal 290, 3512–3526 (2023). 10.1111/febs.16540

74 Tan, C. X. & Eroglu, C. Cell adhesion molecules regulating astrocyte-neuron interactions. Curr Opin Neurobiol 69, 170–177 (2021). 10.1016/j.conb.2021.03.015

75 Araç, D. et al. Structures of neuroligin-1 and the neuroligin-1/neurexin-1 beta complex reveal specific protein-protein and protein-Ca2+ interactions. Neuron 56, 992–1003 (2007). 10.1016/j.neuron.2007.12.002

76 Yamagata, A. et al. Structural insights into modulation and selectivity of transsynaptic neurexin-LRRTM interaction. Nature communications 9, 3964 (2018). 10.1038/s41467-018-06333-8

77 Liouta, K. et al. LRRTM2 controls presynapse nano-organization and AMPA receptor sub-positioning through Neurexin-binding interface. Nature communications 15, 8807 (2024). 10.1038/s41467-024-53090-y

78 Ling, E. et al. A concerted neuron-astrocyte program declines in ageing and schizophrenia. Nature 627, 604–611 (2024). 10.1038/s41586-024-07109-5

79 Lewis, D. A. The human brain revisited: opportunities and challenges in postmortem studies of psychiatric disorders. Neuropsychopharmacology 26, 143–154 (2002). 10.1016/s0893-133x(01)00393-1

80 McCullumsmith, R. E., Hammond, J. H., Shan, D. & Meador-Woodruff, J. H. Postmortem brain: an underutilized substrate for studying severe mental illness. Neuropsychopharmacology 39, 65–87 (2014). 10.1038/npp.2013.239

81 Huitinga, I. & Webster, M. J. Brain Banking. (Elsevier, 2018).

82 Qu, L., Akbergenova, Y., Hu, Y. & Schikorski, T. Synapse-to-synapse variation in mean synaptic vesicle size and its relationship with synaptic morphology and function. J Comp Neurol 514, 343–352 (2009). 10.1002/cne.22007

83 Kirchhausen, T., Owen, D. & Harrison, S. C. Molecular structure, function, and dynamics of clathrin-mediated membrane traffic. Cold Spring Harb Perspect Biol 6, a016725 (2014). 10.1101/cshperspect.a016725

84 Droz, B., Rambourg, A. & Koenig, H. L. The smooth endoplasmic reticulum: structure and role in the renewal of axonal membrane and synaptic vesicles by fast axonal transport. Brain Res 93, 1–13 (1975). 10.1016/0006-8993(75)90282-6

85 Quon, E. & Beh, C. T. Membrane Contact Sites: Complex Zones for Membrane Association and Lipid Exchange. Lipid Insights 8, 55–63 (2015). 10.4137/lpi.S37190

86 Bezprozvanny, I. & Kavalali, E. T. Presynaptic endoplasmic reticulum and neurotransmission. Cell calcium 85, 102133 (2020). 10.1016/j.ceca.2019.102133

87 Drummond, G. B., Paterson, D. J. & McGrath, J. C. ARRIVE: new guidelines for reporting animal research. Experimental physiology 95, 841 (2010). 10.1113/expphysiol.2010.053785

88 Glantz, L. A. & Lewis, D. A. Decreased dendritic spine density on prefrontal cortical pyramidal neurons in schizophrenia. Arch. Gen. Psychiatry 57, 65–73 (2000). 10.1001/archpsyc.57.1.65

89 Glausier, J. R., Kelly, M. A., Salem, S., Chen, K. & Lewis, D. A. Proxy measures of premortem cognitive aptitude in postmortem subjects with schizophrenia. Psychol. Med. 50, 507–514 (2020). 10.1017/s0033291719000382

90 Tegunov, D. & Cramer, P. Real-time cryo-electron microscopy data preprocessing with Warp. Nat Methods 16, 1146–1152 (2019). 10.1038/s41592-019-0580-y

91 Lamm, L. et al. MemBrain v2: an end-to-end tool for the analysis of membranes in cryo-electron tomography. bioRxiv, 2024.2001.2005.574336 (2024). 10.1101/2024.01.05.574336

92 Zheng, S. Q. et al. MotionCor2: anisotropic correction of beam-induced motion for improved cryo-electron microscopy. Nat Methods 14, 331–332 (2017). 10.1038/nmeth.4193

93 Rohou, A. & Grigorieff, N. CTFFIND4: Fast and accurate defocus estimation from electron micrographs. J Struct Biol 192, 216–221 (2015). 10.1016/j.jsb.2015.08.008

94 Burt, A. et al. An image processing pipeline for electron cryo-tomography in RELION-5. FEBS Open Bio 14, 1788–1804 (2024). 10.1002/2211-5463.13873

95 Warshamanage, R., Yamashita, K. & Murshudov, G. N. EMDA: A Python package for Electron Microscopy Data Analysis. J Struct Biol 214, 107826 (2022). 10.1016/j.jsb.2021.107826

96 Pettersen, E. F. et al. UCSF Chimera--a visualization system for exploratory research and analysis. J Comput Chem 25, 1605–1612 (2004). 10.1002/jcc.20084

97 González-Burgos, G. et al. Distinct Properties of Layer 3 Pyramidal Neurons from Prefrontal and Parietal Areas of the Monkey Neocortex. J Neurosci 39, 7277–7290 (2019). 10.1523/jneurosci.1210-19.2019

98 Miyamae, T., Chen, K., Lewis, D. A. & Gonzalez Burgos, G. Distinct physiological maturation of parvalbumin-positive neuron subtypes in mouse prefrontal cortex Journal of Neuroscience 37, 4883–4902 (2017).

99 Rothman, J. S. & Silver, R. A. NeuroMatic: An Integrated Open-Source Software Toolkit for Acquisition, Analysis and Simulation of Electrophysiological Data. Front Neuroinform 12, 14 (2018). 10.3389/fninf.2018.00014

100 Kudoh, S. N. & Taguchi, T. A simple exploratory algorithm for the accurate and fast detection of spontaneous synaptic events. Biosens Bioelectron 17, 773–782 (2002). 10.1016/s0956-5663(02)00053-2

